# Deep Learning in Spatial Transcriptomics: Learning From the Next Next-Generation Sequencing

**DOI:** 10.1101/2022.02.28.482392

**Authors:** A. Ali Heydari, Suzanne S. Sindi

**Affiliations:** Department of Applied Mathematics, University of California, Merced; Health Sciences Research Institute, University of California, Merced

## Abstract

Spatial transcriptomics (ST) technologies are rapidly becoming the extension of single-cell RNA sequencing (scRNAseq), holding the potential of profiling gene expression at a single-cell resolution while maintaining cellular compositions within a tissue. Having both expression profiles and tissue organization enables researchers to better understand cellular interactions and heterogeneity, providing insight into complex biological processes that would not be possible with traditional sequencing technologies. The data generated by ST technologies are inherently noisy, high-dimensional, sparse, and multi-modal (including histological images, count matrices, etc.), thus requiring specialized computational tools for accurate and robust analysis. However, many ST studies currently utilize traditional scRNAseq tools, which are inadequate for analyzing complex ST datasets. On the other hand, many of the existing ST-specific methods are built upon traditional statistical or machine learning frameworks, which have shown to be sub-optimal in many applications due to the scale, multi-modality, and limitations of spatially-resolved data (such as spatial resolution, sensitivity and gene coverage). Given these intricacies, researchers have developed deep learning (DL)-based models to alleviate ST-specific challenges. These methods include new state-of-the-art models in alignment, spatial reconstruction, and spatial clustering among others. However, deep-learning models for ST analysis are nascent and remain largely underexplored. In this review, we provide an overview of existing state-of-the-art tools for analyzing spatially-resolved transcriptomics, while delving deeper into the DL-based approaches. We discuss the new frontiers and the open questions in this field and highlight the domains in which we anticipate transformational DL applications.

## I. INTRODUCTION

Although multicellular organisms contain a common genome within their cells, the morphology and gene expression patterns of cells are largely distinct and dynamic. These differences arise from internal gene regulatory systems and external environmental signals. Cells proliferate, differentiate and function in tissues while sending and receiving signals from their surroundings. These environmental factors cause cell fate to be highly dependent on the environment in which it exists. Therefore, monitoring a cell’s behavior in the residing tissue is crucial to understanding cell function, as well as its past and future fate^1^.

Advancements in single-cell sequencing have transformed the genomics and bioinformatics fields. The advent of singlecell RNA sequencing (scRNAseq) has enabled researchers to profile gene expression levels of various tissues and organs, allowing them to create comprehensive atlases in different species^2–6^. Moreover, scRNAseq enables the detection of distinct subpopulations present within a tissue; which has been paramount in discovering new biological processes, the inner workings of diseases, and effectiveness of treatments^7–14^. However, high-throughput sequencing of solid tissues requires tissue dissociation, resulting in the loss of spatial information^15,16^. To fully understand cellular interactions, data on tissue morphology and spatial information is needed, which scRNAseq alone can not provide. The placement of cells within a tissue are crucial from the developmental stages (*e.g*. asymmetric cell fate of mother and daughter cells^17^) and beyond cell differentiation (such as cellular functions, response to stimuli and tissue homeostasis^18^). These limitations would be alleviated by technologies that could preserve spatial information while measuring gene expression at the singlecell level.

Spatial Transcriptomics (ST) provide an *unbiased* view of tissue organization crucial in understanding cell fate, delineating heterogeneity, and other applications^19^. However, many current ST technologies suffer from lower sensitivities as compared to scRNAseq, while lacking the single-cell resolution that scRNAseq provides^20^. Targeted *in situ* technologies have tried to solve the issue of resolution and sensitivity, but are limited in gene throughput and often require *a priori* knowledge of target genes^20^. More specifically, *in situ* technologies (such as *in situ* sequencing^21^, single-molecule fluorescence *in situ* hybridization (smFISH)^22–24^, targeted expansion sequencing^25^, cyclic-ouroboros smFISH (osmFISH)^26^, multiplexed error-robust fluorescence *in situ* hybridization (MERFISH)^27^, sequential FISH (seqFISH+)^28^, and spatially resolved transcript amplicon readout mapping (STARmap)^29^), are typically limited to pre-selected genes that are on the order of hundreds, with the accuracy potentially dropping as more probes are added^29^. We will refer to these methods as *imagebased* techniques.

On the other hand, Next Generation Sequencing (NGS)-based technologies (such as 10x Genomics’ Visium and its predecessor^30,31^, Slide-Seq^32^, HDST^33^) barcode entire transcriptomes but have limited capture rates, and resolutions that are larger than a single cell^34^ (50 *μ*m - 100 *μ*m for Visium and 10 *μ*m for Slide-Seq). Moreover, unlike image-based technologies, NGS-based methods allow for unbiased profiling of large tissue sections without necessitating a set of target genes^35,36^. However, NGS-based technologies do not have single-cell resolution, requiring cellular features to be inferred or related to the histological scale using computational approaches. Many current algorithms use traditional statistical or medical image processing frameworks that require human supervision^34,37,38^, which is not ideal for large-scale analyses. Additionally, many algorithms are not generalizable across different sequencing platforms, which limit their utility and restrict multiomics integration efforts.

**FIG. 1.**
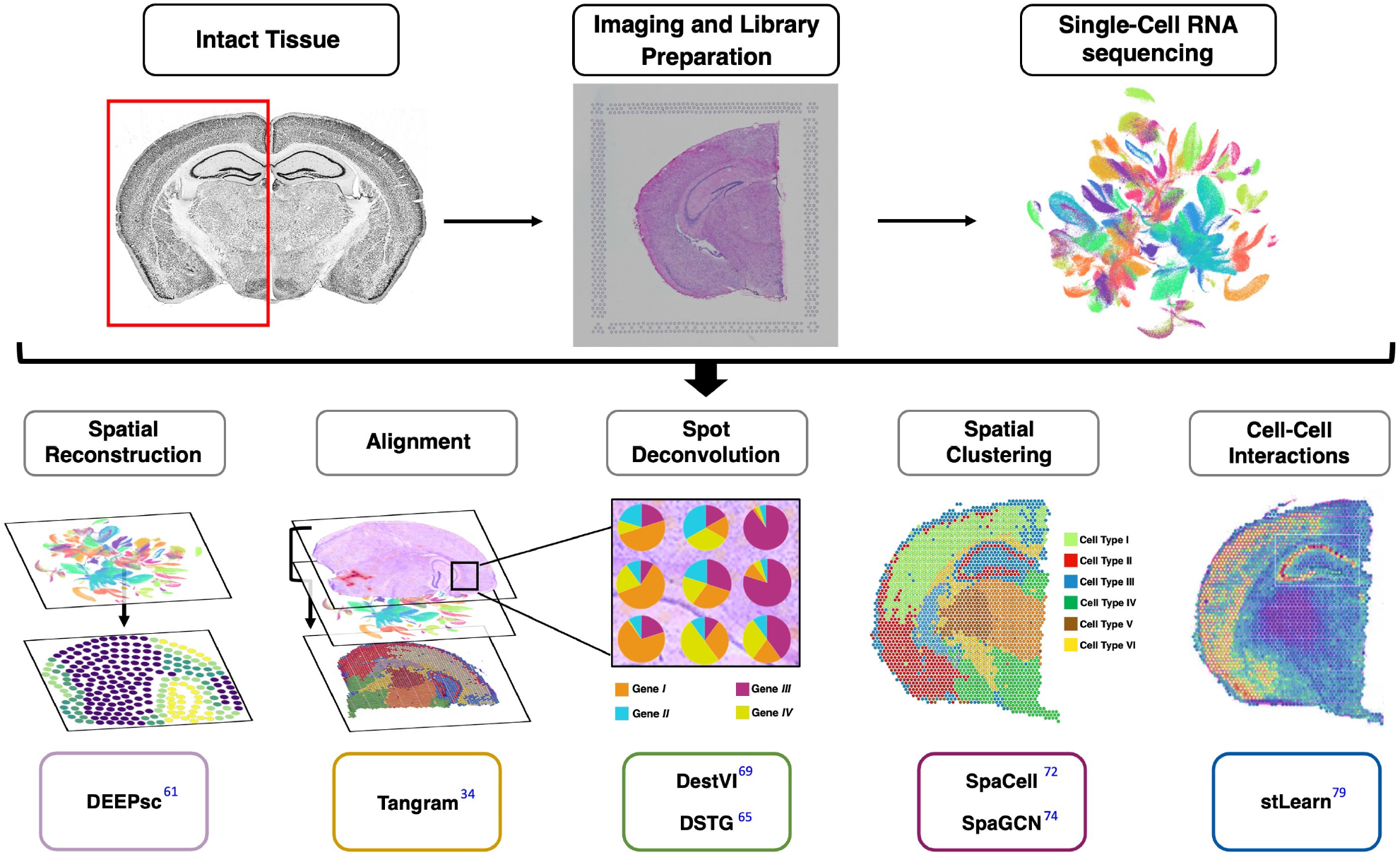
An overview of Deep Learning methods for spatial transcriptomics presented in this review. We provide a more comprehensive list of the state-of-the-art methods for spatial transcriptomics in Table I.

Deep Learning (DL) methods can use raw data to extract useful representations (or information) needed for performing a task, such as classification or detection^39^. This quality makes this class of Machine Learning (ML) algorithms ideal for applications where the available data is large, higher-dimensional, and noisy, such as single-cell omics. DL models have been extensively used in scRNAseq studies (*e.g*. preprocessing^40,41^, clustering^42,43^, cell-type identification^44,45^ and data augmentation^46,47^), and have shown to significantly improve upon traditional methods^10^, suggesting the potential of such methods in ST analysis. Moreover, DL models can leverage multiple data sources, such as images and text data, to learn a set of tasks^48^. Given that spatially-resolved transcriptomics are inherently multimodal (i.e. they consist of images and gene expression count data) and that downstream analysis consist of multiple tasks (*e.g*. clustering and cell-type detection), researchers have sought to develop ST-specific DL algorithms.

Spatially-resolved transcriptomics have been utilized to unravel complex biological processes in many diseases (*e.g*. COVID-19^49,50^, arthritis^51,52^, cancer^31,33,53–55^, Alzheimer’s^56^, diabetes^57,58^, etc.). Continuous improvements and commercialization of ST technologies (such as 10x’s Visium) are resulting in wider use across individual labs. Therefore, scalable and platform-agnostic computational approaches are needed for accurate and robust analysis of ST data. So far, DL methods have shown promising results in handling the scale and multi-modality of spatially-resolved transcriptomics; however, DL-based models in this space remain nascent. Similar to scRNAseq analysis, we anticipate a suite of DL models to be developed in the near future to address many of the pressing challenges in spatial omics field. This review aims to provide an overview of the current state-of-the-art (SOTA) DL models developed for ST analysis. Due to the potentials and accessibility of NGS-based ST technologies, we primarily focus on methods and techniques developed for these technologies.

The remainder of this manuscript is organized as follows: We provide an overview of common scRNAseq and ST technologies in Section II, followed by a general description of common DL architectures used for ST analysis in Section III. Section IV is dedicated to the current DL methods developed for analyzing spatially-resolved transcriptomics. We conclude, in Section V, by discussing our outlook on the current challenges and future research directions in ST domain. Table I provides the reader with a list of current SOTA methods for ST analysis. Given the pace of advancements in this field, the authors have compiled an online list of current DL methods for ST analysis on a dedicated repository (https://github.com/SindiLab/Deep-Learning-in-Spatial-Transcriptomics-Analysis), which will be maintained and continuously updated.

**TABLE I:**
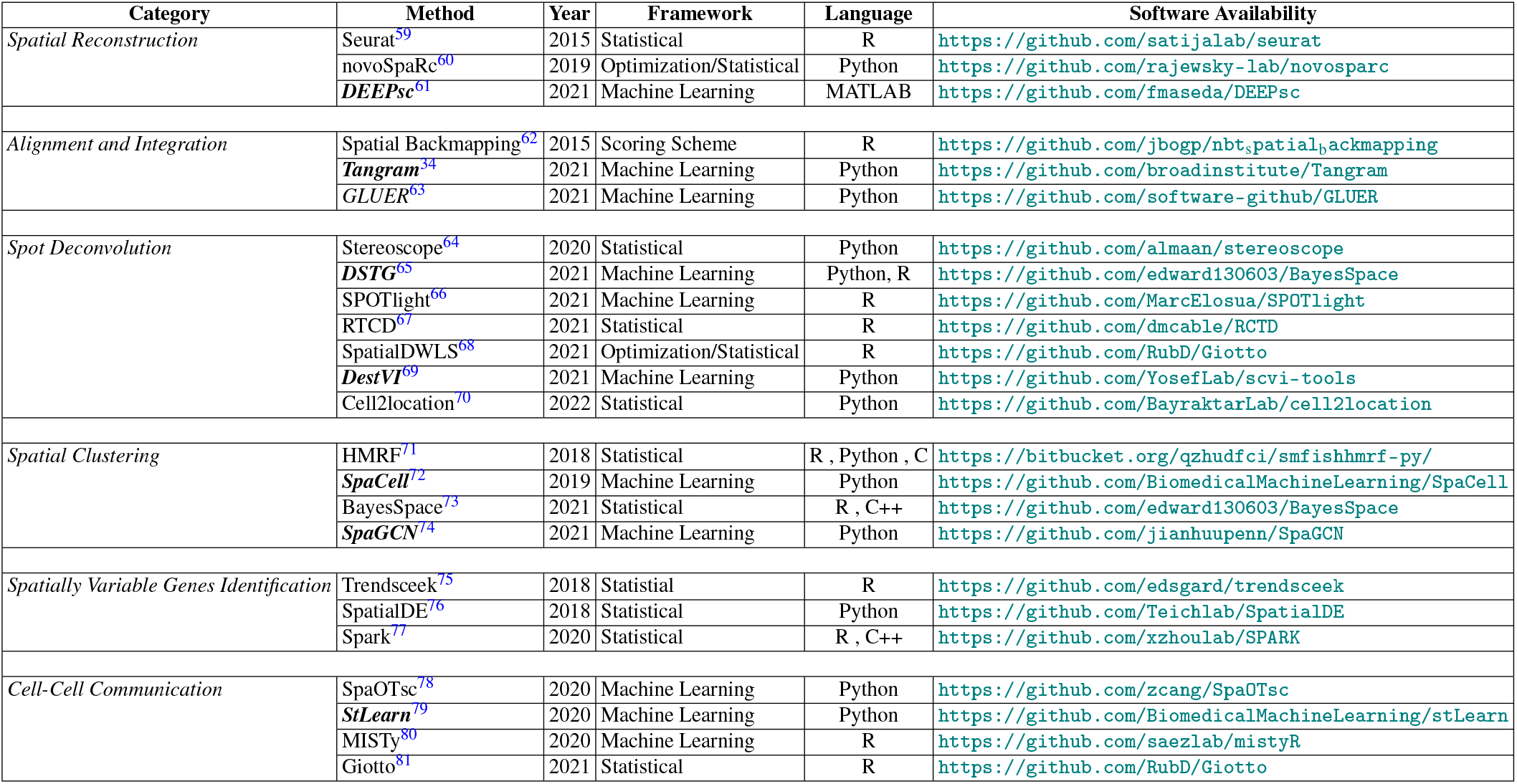
A list of relevant methods for the analysis of spatial transcriptomics data. The italicized boldfaced methods are the ones which utilize deep learning (or elements closely aligned). We review these methods in depth in this paper.

## II. BIOLOGICAL BACKGROUND

### A. Single-Cell RNA Sequencing (scRNAseq)

RNA sequencing (RNA-seq) provides comprehensive insights on cellular processes (such as identifying genes that are upregulated or downregulated, etc.). However, traditional bulk RNA-seq is limited to revealing the average expression from a collection of cells, and not disambiguation single-cell behavior. Thus, it is difficult to delineate cellular heterogeneity with traditional RNA-seq, which is a disadvantage since cellular heterogeneity has been shown to play a crucial role in understanding many diseases^82^. Therefore, researchers have turned to single-cell RNA-seq (scRNAseq) in order to identify cellular heterogeneity within tissues. ScRNAseq technologies have been instrumental in the study of key biological processes in many diseases, such as cancer^83^, Alzheimer’s^84^, cardiovascular diseases^85^, etcetera (see^82^ for more details). RNA sequencing of cells at a single-cell resolution, scRNAseq, generally consists of four stages:

i. **Isolation of Single-Cells and Lysing**: Cells are selected through laser microdirection, fluorescence-activated cell sorting (FACS), microfluidic/microplate Technology (MT) or a combination of these methods^86^, with MT being highly complementary to NGS-based technologies^87^. MT encapsulates each single-cell into an independent microdroplet containing unique molecular identifiers (UMI), lysis buffer for cell lysis (to increase the capturing of as many RNA molecules as possible), oligonucleotide primers, and seoxynucleotide triphosphates (dNTPs) in addition to the cells themselves. Due to MT’s higher isolation capacity, thousands of cells can be simultaneously tagged and analyzed, which is beneficial for large-scale scRNAseq studies.
ii. **Reverse Transcription**: One challenge in RNA sequencing is that RNA can not be directly sequenced from cells, and thus RNA must first be converted to complementary DNA (cDNA)^88^. Although dist technologies employ different techniques, the reverse transcription phase generally involves capturing mRNA using poly[T] sequence primers that bind to mRNA ploy[A] tail prior to cDNA conversion. Based on the sequencing platform, other nucleotide sequences are added to the reverse-transcription; for example in NGS protocols, UMIs are added to tag unique mRNA molecules so that it could be trace back their originating cells, enabling the combination of different cells for sequencing.
iii. **cDNA Amplification**: Given that RNA can not be directly sequenced from cells, single-stranded RNAs must first be reverse-transcribed to cDNA. However, due to the small amount of mRNA in cells, limited cDNA is produced which is not optimal for sequencing. Therefore, the limited quantity of cDNA must be amplified prior to library preparation and sequencing^89^. The amplification is often done by either PCR (exponential amplification process with its efficiency being sequence dependent) or IVT (a linear amplification method which requires an additional round of reverse transcription of the amplified RNA) before sequencing^88,90^. The final cDNA library consists of adaptor-ligated sequencing library attached to each end.
iv. **Sequencing Library Construction**: Finally, every cell’s tagged and amplified cDNA is combined for library preparation and sequencing similar to bulk RNA sequencing methods, followed by computational pipelines for processing and analysis.^91^.

Fig. 2(A) illustrates an example of the workflow for scRNAseq. For more details of each stage and various scRNAseq workflows, we refer the reader to references^90,92–94^.

**FIG. 2.**
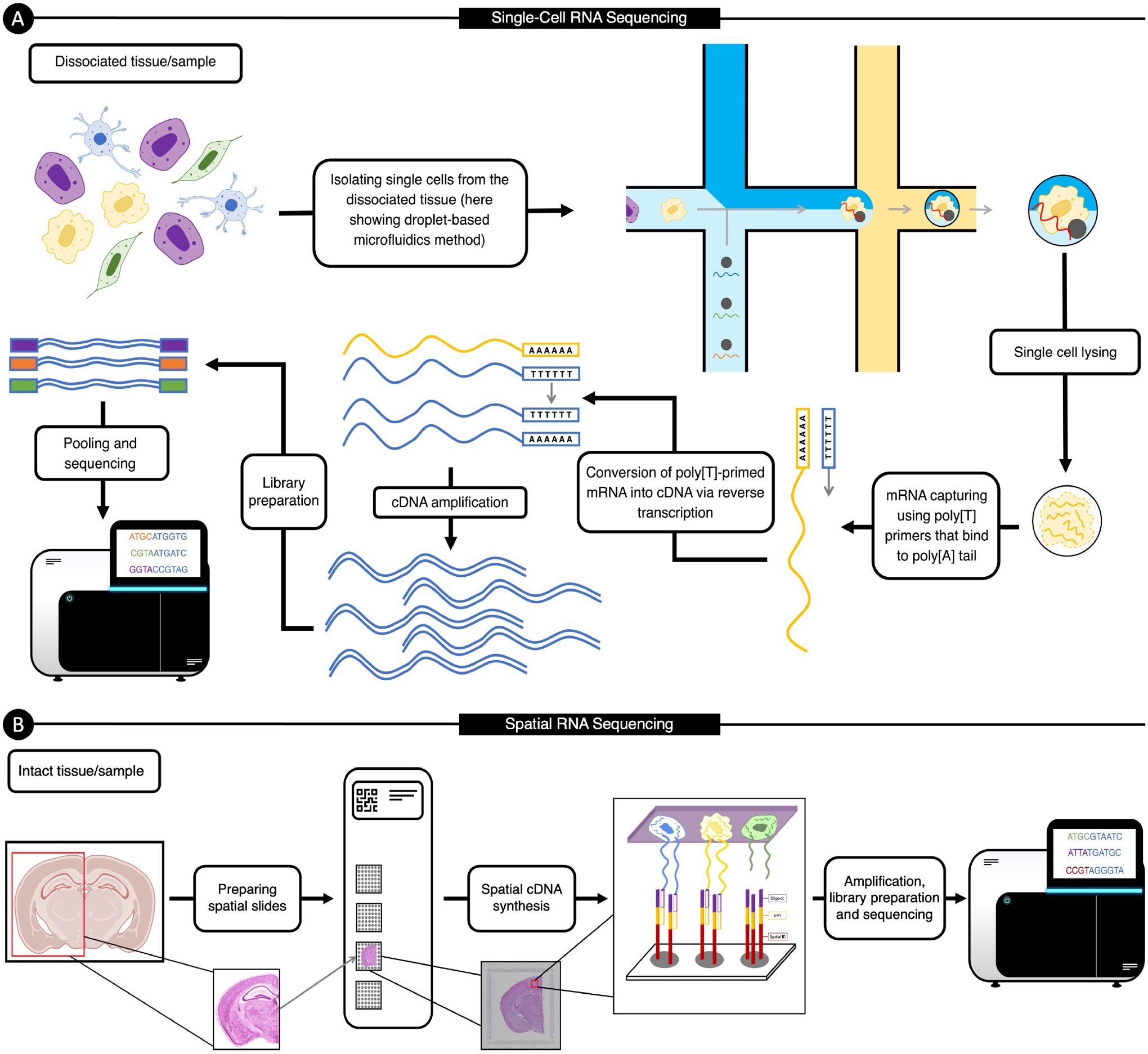
Example Single-Cell RNA Sequencing and Spatial Transcriptomics Workflows. **(A)** A general overview of the single-cell RNA sequencing workflow (which we describe in section II A). **(B)** A visualization of the steps for next-generation sequencing based spatial transcriptomics (described in Section II B).

### B. Spatial Transcriptomics Technologies

More recently, technologies that profile gene expression while retaining spatial information have emerged. These technologies are collectively known as *spatial transcriptomics* (ST). The various ST technologies provide different advantages and are chosen based on experimental factors such as size of tissue to be assayed, the number of genes to be probed, *a priori* knowledge of taget genes, cost, etcetera. In general, ST technologies can be divided into two broad categories: imaging-based and next generation sequencing (NSG)-based technologies. In this section, we provide an overview of popular techniques, with more emphasis on NGS-based approaches. For a more comprehensive and technical reviews of ST technologies, we refer the reader to Asp *et al*.^95^ and Rao *et al*.^20^.

#### 1. Imaging-Based Technologies

Imaging-based technologies are be broadly subdivided into *in situ* hybridization (ISH)-based, *in situ* sequencing (ISS)-based methods, or methods that borrow elements from both of these approaches. Unlike RNAseq methods described above, ISH and most ISS-based techniques require labeled probes. This means that the target genes must be known in advance and, moreover, the number of genes that can be measured is limited^20^.

##### *In Situ* Hybridization (ISH)-based Approaches

ISH-based methods aim to detect the absence or presence of target RNA (or DNA) sequences while localizing the information of the desired sequences to specific cells or chromosomal sites^96,97^.

ISH-based techniques use labeled probes (usually made with DNA or RNA) which bind to desired sequences in fixed cells or tissue, therefore detecting the desired sequence through the hybridization of a complementary probe. The hybridized probes are then visualized through isotopic and nonisotopic (fluorescent and nonfluorescent) approaches^97^. The ISH-based techniques have been limited by the number of distinguishable transcripts, however, recent innovations have resulted in ample multiplexing capabilities^20^.

##### *In Situ* Sequencing (ISS)-based Approaches

ISS-based approaches aim to sequence the RNA content of a cell *in situ* using DNA balls that amplify the RNA signals: RNA is first reverse transcribed to cDNA, followed by circular amplification (to increase the number of transcripts) and sequencing^98^. Although the transcript can be localized at subcellular resolution, micrometer- or nanometer-sized DNA balls are often used to amplify the signals to reach sufficient signal for imaging^95^. Initially, the first ISS-based method^99^ used targeted padlock probes (a single-stranded DNA molecule containing regions complementary to the target cDNA) followed by sequence by-ligation^21^ to detect desired genes. This method provided a subcellular resolution and an ability to detect singlenucleotide variants (SNVs). This ISS protocol is targeted and yields a detection efficiency of approximately 30%^100^. Several ISS protocols have built upon this approach to mitigate the number of cells that can be discriminated simultaneously, as well as to improve certain experimental aspect of the protocol. For example, a recently developed method, *barcode insitu targeted sequencing* (BaristaSeq)^101^, uses sequencing-by-synthesis and has led to increased read lengths, enabled higher throughput and cellular barcoding with improved detection efficiency compared to the initial ISS approach^101^. Another ISS-based technique is Spatially Resolved Transcript Amplicon Readout Mapping (STARmap)^29^ that reduces noise and avoids the cDNA conversion complications by utilizing improved padlock-probe and primer design; STARmap adds a second primer to target the site next to the padlock probe in order to circumvent the reverse transcription step. STARmap also uses advanced hydrogel chemistry and takes advantage of an error-robust sequencing-by-ligation method, resulting in detection efficiency that is comparable to scRNAseq methods (around 40%)^29,95^. Although most ISS approaches (including the ones mentioned here) are targeted, ISS-based methods could also be untargeted^25,102^ but this typically leads to much lower sensitivity (around 0.005%) and molecular crowding, affecting the rolling-circle amplification bias^102,103^.

In imaging-based approaches, the generated image is segmented and processed to produce a cell-level gene-expression matrix. The gene-expression matrix is generated through processing the generated image(s), which can be done manually or automatically. However, given the biased and laborious nature of manual segmentation, there has been a shift towards designing general and automated techniques^104^. The accurate and general automation of this process still remains a challenge, therefore motivating the application of recent machine learning and computer vision approaches to this field^104–106^, which have shown improvements compared to the traditional methods^107^. Although this manuscript focuses on methods for NGS-based technologies, many of those techniques (including ones in Section IV C and IV F) can be extended to image-based technologies as well.

### C. Next Generation Sequencing (NGS)-Based Technologies

Due to the unbiased capture of mRNA, NGS-based technologies can shed light on the known and unknown morphological features using only the molecular characterization of tissues^1^. This unbiased and untargeted nature of NGS technologies makes them ideal for studying and exploring new systems^20^, a major advantage compared to most image-based technologies which require target genes *a priori*. While NGS-based approaches differ in the specifics of the protocols, they all build on the idea of adding spatial barcodes before library preparation, which are then used to map transcripts back to the appropriate positions (known as spots or voxels). An example workflow of NGS-based spatial sequencing is depicted in Fig. 2(B). In the following subsections, we provide a general overview of the four most common spatial transcriptomics technologies. For a more complete review of these technologies, we refer the readers to references^20,95^.

Ståhl *et al*.^31^ were the first to successfully demonstrate the feasibility of using NGS for spatial transcriptomics (this initial approach is often referred to *Spatial Transcriptomics*). Their innovation was to add spatial barcodes prior to library preparation, enabling the mapping of expressions to appropriate spatial spots. More specifically, Ståhl *et al*. positioned oligo(dT) probes and unique spatial barcodes as microarrays of spots on the surface of slides. Next, fresh frozen tissue slices were placed on the microarray and processed to release mRNA (using enzymatic permeabilization), which then hybridized with the probes on the surface of the slides. This approach consists of (i) collecting histological imaging (using standard fixation and staining techniques, including hematoxylin and eosin (HE) staining) for investigating morphological characteristics and (ii) sequencing spatially barcoded cDNA to profile gene expressions. In the initial experiments, each slide consisted of approximately 1000 spots, each of diameter 100 *μm* with 200 *μm* center-to-center distance^1^. This approach provides researchers with an unbiased technique for analyzing large tissue areas without the need for selecting target genes in advance^20,35,108^.

After the initial success of *Spatial Transcriptomics*, 10x Genomics subsequently improved the resolution (shrinking the spot diameters to 55 *μm* with 100 *μm* centre-to-centre distance) and sensitivity (capturing more than 10^4^ transcripts per spot) of the approach, and eventually commercializing it as Visium^30,109^). The development and commercialization of the spatial transcriptomics resulted in relatively rapid adoption across fields, such as cancer biology^110,111^, developmental biology^112,113^, neuroscience^114,115^. The histological imaging and gene expression profiling of Visium are similar to the initial approach: the staining and imaging of the tissues are through traditional staining techniques, including HE staining for visualizing tissue sections using a brightfield microscope and immunofluorescence staining to visualize protein detection in tissue sections through a fluorescent microscope. Visium protocol allows for both fresh frozen (FF) or Formalin-Fixed Paraffin-Embedded (FFPE) tissues. For FF tissues, similar to Ståhl *et al*.^31^, the tissue is permeabilized, allowing the release of mRNA, which hybridizes to the spatially barcoded oligonucleotides present on the spots. The captured mRNA then goes through a reverse transcription process that results in cDNA, which are then barcoded and pooled for generating a library^116^. For FFPE tissues, tissue is permeabilized to release ligated probe pairs from the cells that bind to the spatial barcodes on slide, and the barcoded molecules are pooled for downstream processing to library generation^116^.

Building on the *Spatial Transcriptomics*, Vickovic *et al*.^33^ proposed High-Definition Spatial Transcriptomics (HDST) which improved the resolution to about 2 *μm*. Similar to the other approaches, HDST also employs specific barcodes ligated to beads that are coupled to a spot (prior to lysis), so that expressions are mapped to the tissue image. However, the innovation of HDST include the use of 2 *μm* beads places in hexagonal wells, enabling accurate compartmentalization and grouping of the biological materials in the experiment^33^. Simultaneously, Rodriques *et al*.^32^ introduced SlideSeq which utilizes slides with randomly barcoded beads to capture mRNA, also increasing the resolution (to 10 *μm*) and sensitivity (500 transcripts per 54 bead) spatial-resolved sequencing compared to Ståhl *et al*.^31^. However, SlideSeq placed the barcoded beads in rubber and onto glass slides, as opposed to HDST’s hexogonal beads, and determines the position of each random barcode by *in situ*-indexing^20,32^.

Despite the differences, all NGS-based technologies use spatial barcodes to tag released RNAs, which then go through conventional processes for sequencing similar to scRNAseq. After sequencing, the data is processed to construct the spatial location of each read (using the spatial barcode) and to construct a gene-expression matrix (mapping the reads to the genome to identify the transcript of origin). Given that most technologies have resolutions larger than a single-cell (commonly having expression for 3 to 30 cells in each spot), the data processing and analysis procedures are relatively similar.

## III. MACHINE LEARNING AND DEEP LEARNING BACKGROUND

With the technologies now defined, we next describe common *Machine Learning* (ML) methods used to analyze ST data. in this section, we first provide a discussion of the algorithmic development of ML and *Deep Learning* (DL) models, and then discuss common architectures used for spatially-resolved transcriptomics (and scRNAseq data).

ML refers to a computer algorithm’s ability to acquire knowledge by extracting patterns and features from raw data^117^. All ML algorithms depend on data, which must be available before the methods can be used, and a defined mathematical objective. ML models’ lifecycle consists of two phases, namely *training* and *evaluation*. During training, ML algorithms analyze the data to extract patterns and adjust their internal parameters based on optimizing their objectives (known as *loss function*). In the evaluation (or inference) stage, the trained model makes predictions (or performs the task it was trained to do) on *unseen* data.

There are two main types of ML algorithms: *supervised* and *unsupervised*. An ML algorithm is considered to be *unsupervised* if it utilizes raw inputs without any labels to optimize its objective function (an example would be the K-Means clustering algorithm^118^). Conversely, if an algorithm uses both raw data and the associated labels (or targets) in training, then it is a *supervised* learning algorithm. Supervised learning is the most common form of ML^39^. An example of supervised learning in scRNAseq analysis would be classifying cell subpopulations using prior annotations: this requires a labeled set of cell-types for training (the available annotations), an objective function for calculating learning statistics (“teaching” the model), and testing data for measuring how well the model can predict the cell-type (label) on data it has not seen before (*i.e*. generalizibility of the model). Another common example of supervised learning is regression, where a model predicts continuous values as opposed to outputting labels or categorical values in classification. For supervised tasks, a model is trained on the majority of the data (known as *training set*) and then evaluated on held-out data (*test set*). Depending on the size of our dataset, there can also be a third data split known as a *validation set*, which is used to measure the performance of the model throughout training to determine *early stopping*^119^: Early stopping is when we decide to stop the training of a model because its overfitting (or over optimization) on the training set. Overfitting on training data worsens the generalizability of the model on unseen data, which early stopping aims to avoid^119^. In addition to supervised and unsupervised algorithms, there are also *semi-supervised* learning, where a model uses a mix of both supervised and unsupervised tasks, and *self-supervised*, where the computer algorithm generates new or additional labels to improve its training, or to learn a new task.

Raw experimental data typically contains noise or other unwanted features, which present many challenges for ML algorithms. Therefore, it is often necessary to carefully preprocess data or to rely on domain-specific expertise in order to transform raw data into some internal representation from which ML models can learn^39^. Deep Learning (DL) algorithms, however, aim to use only raw data to automatically extract and construct useful representations required for learning the tasks at hand. In a broad sense, DL models are able to learn from observations through constructing a hierarchy of concepts, where each concept is defined by its relation to simpler concepts. A graph representation of the hierarchy of concepts (and learning) will consist of many layers, with many nodes and edges connecting the vertices, somewhat resembling humans’ neural network. This graph is referred to as an Artificial Neural Network (ANN). ANNs are composed of interconnected nodes (“artificial neurons”) that resemble and mimic our brains’ neuronal functions. An ANN is considered to be a DL model if it consists of many layers–often more than three, hence being called *deep*.

Many tasks that humans perform can be viewed as mappings between sets of inputs and outputs. For example, humans can take a snapshot image of their surroundings (input) and detect the relevant objects (the outputs). DL, and more generally Artificial Intelligence, aims to learn such mappings in order to model human-level intelligence. Mathematically, ANNs are universal function approximators, meaning that, theoretically, they can approximate any (continous) function^120–122^. Cybenko^120^ proved this result for a one-layer neural network with arbitrary number of neurons (nodes) and a sigmoid activation function by showing that such architecture is dense within the space of continuous functions (this result has now been extended to ANNs with multiple layers^121^). While constructing arbitrarily-long single-layer ANNs is not possible, it has been shown that ANNs with many many layers (deeper) generally learn faster and more reliably than ANNs with few wide (many neurons) layers^123^. This has allowed researchers to employ deep networks for learning very complex functions through constructing simple non-linear layers which can transform the representation of each module (starting with the raw input) into a representation at a higher, slightly more abstract level^39^.

DL models’ ability to approximate highly non-linear functions has revolutionized many domains of science, including Computer Vision^124–126^, Natural Language Processing^127–129^ and Bioinformatics^130–133^. DL is becoming increasingly incorporated in many computational pipelines and studies, specially in genomics and bioinformatics, including scRNAseq and spatial transcriptomics analysis. In the following sections, we provide a brief overview of essential deep learning architectures that have been used in spatial transcriptomics and scRNAseq analysis. In Fig. 3, we present illustrations of the architectures discussed in the following sections. Note that for simplicity, we have categorized all Graph Convolution Networks (GCN)^134^ as DL models; this is because (i) GCNs can easily be extended to include more layers (deeper networks), and (ii) lack of other existing methods which incorporate some elements of DL. A more comprehensive description of each architecture can be found in the seminal textbook by Goodfellow *et al*.^117^.

**FIG. 3.**
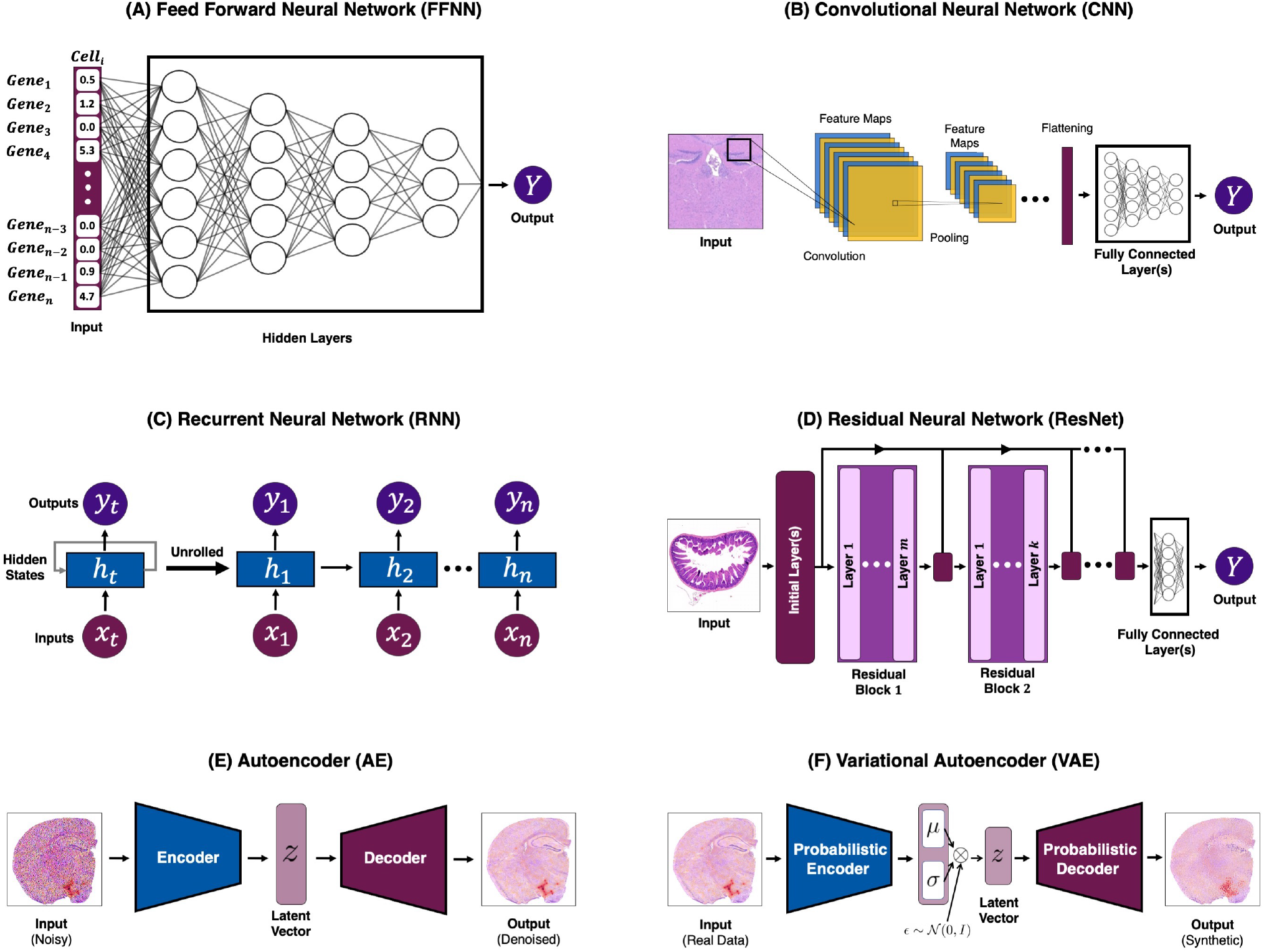
Examples of Deep Learning Architectures. Models depicted in **(A), (B), (C), (D)** are examples of supervised learning, and networks shown in **(E), (F)** are unsupervised. **(A)** An example of an FFNN architecture with gene expression count as its input. **(B)** An example of CNN architecture, where the model passes the inputs through the three stages of a CNN (with non-linear activation not depicted) to extract features. Then, outputs are flattened and fed into a fully connected layer (or layers). **(C)** The general training flow of an RNN, with the unrolled version showing the timestep-dependent inputs, hidden state, and outputs. The inputs to RNNs need to have a sequential structure (*e.g*. time-series data). **(D)** An illustration of a ResNet. In traditional ResNets, there are identity mappings (or skip connections) that pass the input of a residual block to its output (often through addition). **(E)** Here we show the general architecture of a *trained* denoising AE in inference stage, with a noisy histology slide as its input, yielding a denoised version of the input image. **(6)** A depiction of a traditional VAE in inference stage. VAE’s aim to generate synthetic data that closely resemble the original input. This is done through regularizing the latent space of an AE with the use of a probabilistic encoder and decoder.

### A. Feed Forward Neural Network (FFNN)

FFNNs, the quintessential example of Artificial Neural Networks (ANNs), aim to approximate a function mapping a set of inputs to their corresponding targets (see Fig. 3(A)). More specifically, given an input 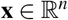 and a target 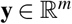, where *n*, 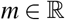, FFNNs aim to *learn the optimal parameters **θ** such that* **y** = *f*(**x**; ***θ***). FFNNs are the building blocks of many more advanced architectures (*e.g*. convolutional neural networks), and therefore, of paramount importance in the field of ML^117^. As mentioned previously, ANNs are universal function approximators, and they represent a directed acyclic graph of function compositions hierarchy within the network. Each layer of a FFNN, *f*^(*i*)^(**x**; ***θ***) (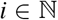 being the *i*-th layer), is often a simple linear function: For example, we can have a linear function for outputting 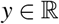 of the form Eq. (1), with weight parameters 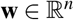 and a bias 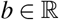:

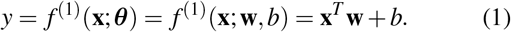

However, a model composed of *only* linear functions can *only* approximate linear mappings. As such, we must consider *non-linear activation* functions to increase model capacity, enabling the approximation of complex non-linear functions. In the simplest case, Neural networks (NNs) use an affine transform (controlled by learned parameters) followed by a non-linear activation function, which, theoretically, enables them to approximate any non-linear function^135^. Moreover, we could compose many of such non-linear transformations to avoid infinitely wide-neural networks when approximating complex function. However, in this context, finding a set of optimal functions 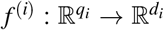 (*q_i_*, 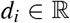) is a practically impossible task. As such, we restrict the class of function that we use for *f*^(*i*)^ to the following form in Eq. (2):

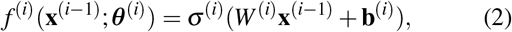

where superscript *i* enumerates the layers, ***σ***(·) is a nonlinear activation function (usually a Rectified Linear Unit^136^), 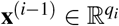 denotes the output of the layer (*i* – 1) (with **x**^(0)^ indicating the input data), weights 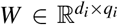 and biases 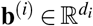. Note that because of the dimensionality of the mapping, 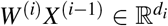 and we must have a vector of biases 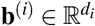). FFNNs are composed of such functions in chains; to illustrate, consider a three-layer neural network:

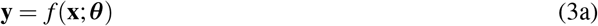

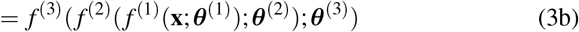

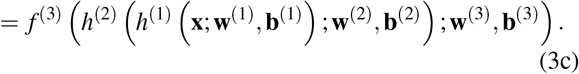

with *h* representing the *hidden states or hidden layers*.

FFNNs find the optimal contribution of each parameter (*i.e*. weights and biases) by minimizing a desired objective. The goal is to generalize the task to data the model has never seen before (testing data). Although the non-linearity increases the capacity of FFNNs, it causes most objective functions to become non-convex. In contrast to convex optimization, non-convex loss functions do not have global convergence guarantees, and are sensitive to initial starting point (parameters of the network)^137^. Therefore, such optimization is often done through stochastic gradient descent (or some variant of it). Moreover, given the sensitivity to initial values, weights are typically chosen to be small random values, with biases initialized to zero or small positive values^117,138,139^.

Almost all neural networks use iterative gradient descent (GD)-based optimizers to train. GD has three main variants, which differ in the amount of data utilized to calculate the gradients for updating the parameters. The classic GD variant, referred to as *batch* GD, uses all data points to make the updates to the parameters in *one iteration*. However, this approach is generally not feasible, since the amount of data required for training DL models almost never fits in memory^140^. The second variant of GD is *stochastic gradient descent* (SGD) where the parameters are updated for every training datum. Computationally, it has been shown that the noise in SGD accelerates its convergence compared to batch GD, but SGD also has the possibility of overshooting, specially for highly non-convex optimization functions^140,141^. The third variant, and the most frequently used one for deep learning, is mini-batch GD which updates the parameters for every batch of training data–if batch size is one then this variant is just SGD, and if batch size is the entire dataset then it is equivalent to batch GD. Conventionally, optimization of NNs is done through gradient descent performed backwards in the network, which itself consists of two components: a numerical algorithm for efficient computation of the chain rule for derivatives (backpropagation^142^) and a GD-based optimizer (*e.g*., Adam^143^ or AdaGrad^144^). The optimizer is an algorithm that performs gradient descent, while backpropagation is an algorithm for computing the expression for gradients during the backward pass of the model.

### B. Convolutional Neural Network (CNN)

Learning from images, such as detecting edges and identifying objects, has been of interest for some time in computer science^145^. Images contain a lot of information, however, only a small amount of that information is often relevant to the task at hand. For example, an image of a stained tissue contains both important information, namely the tissue itself, and irrelevant pixels, such as the background. Prior to DL, researchers would hand-design a feature extractor to learn relevant information from the input. Much of the work had focused on the appropriate feature extractors for desired tasks (*e.g*. see the seminal work by Marr and Hildreth^146^). However, a main goal in ML is to extract features from raw inputs without hand-tuned kernels for feature extraction. CNNs^147,148^ are a specialized subset of ANNs that use the convolution operation (in at least one of their layers) to learn appropriate kernels for extracting important feature beneficial to the task at hand. Mathematically, convolution between two functions *f* and *w* is defined as a commutative operation shown in Eq. (4)

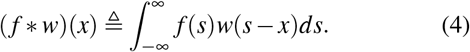

Using our notation, we intuitively view convolution as the area under *f*(*s*) weighted by *w*(–*s*) and shifted by *x*. In most applications, discrete functions are used. As an example, assume we have a 2D kernel *K* that can detect edges in a 2D image *I* with dimension *m* × *n*. Since *I* is discrete, we can use the discrete form of Eq. (4) for convolution of *I* and *K* over all pixels:

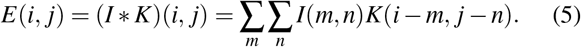

However, since there is less variation in the valid range of *m*, *n* (the dimensions of the image) and the operation is commutative, most algorithms implement Eq. (5) equivalently:

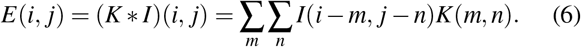

Typical CNNs consist of a sequence of layers (usually three) which include a layer performing convolution, hence called a *convolutional layer* (affine transform), a detector stage (non-linear transformation), and a pooling layer. The learning unit of a convolutional layer is called a *filter or kernel*. Each convolutional filter is a matrix, typically of small dimensions (*e.g*. 3×3 pixels), composed of a set of weights that acts as an object detector, with the weights being continuously calibrated during the learning process. CNNs’ objective is to learn an optimal set of filters (weights) which can detected the needed features for specific tasks (*e.g*. image classification). The result of convolution between the input data and the filter’s weights is often referred to as a *feature map* (as shown in Fig. 3(B)). Once a feature map is available, each value of this map is passed through a non-linearity (*e.g*. ReLU). The output of a convolutional layer consists of as many stacked feature maps as the number of filters present within the layer.

There are two key ideas behind the design of CNNs: First, in data with grid-like topology, local neighbors have highly correlated information. Second, equivariance to translation can be obtained if units at different locations *share weights*. In other words, sharing parameters in CNNs enabled the detection of features regardless of the locations that they appear in. An example of this would be detecting a car. In a dataset, a car could appear at any position in a 2D image, but the network should be able to detect it regardless of the specific coordinates^145^. These design choices provide CNNs with three main benefits compared to other ANNs: (i) sparse interactions, (ii) shared weights, and (iii) equivariant representations^147^.

Another way of achieving equivariance to translation is to utilize pooling layers. Pooling decreases the dimension of learned representations, and makes the model insensitive to small shifts and distortions^39^. In the pooling layers, we use the outputs of the detector stage (at certain locations) to calculate a summary statistic for a rectangular window of values (*e.g*. calculating the mean of a 3×3 patch). There are many pooling operations, with common choices being max-pooling (taking the maximum value of a rectangular neighborhood), mean-pooling (taking the average), and L2 norm (taking the norm). In all cases, rectangular patches from one or several feature maps are inputted to the pooling layer, where semantically similar features are merged into one. CNNs typically have an ensemble of stacked convolution layers, non-linearity, and pooling layers, followed by fully connected layers that produce the final output of the network. The backpropagation of gradients through CNNs is analogous to FFNNs, enabling the model to learn an optimal set of filters for the task(s) at hand. CNNs have been effectively used in many applications in computer vision and time-series analysis, and are being increasing utilized for analysis of ST data, since spatial-omics are multi-modal, with one of the modalities being images (as we discuss in Section IV).

### C. Recurrent Neural Network (RNN)

Just as CNNs are specialized to process data with a grid-like toplogy, RNNs’^149^ special characteristics make them ideal for processing sequential data *X* = {**x**^(1)^, **x**^(2)^,…, **x**^(*n*)^}, where **x**^(*i*)^ denotes the *i*-th element in the ordered sequence *X*. Examples of such sequence-like structure include times series and natural language. RNNs process sequential inputs one at a time and implicitly maintain a history of previous elements of the input sequence. We present an illustration of the conventional RNN architecture in Fig. 3(C). Similar to FFNNs or CNNs, RNNs can be composed of many layers, with each layer depending on the previous hidden state, *h*^(*t*–1)^, and a shared set of parameters, ***θ***. A deep RNN with *n* hidden states can be expressed as follows:

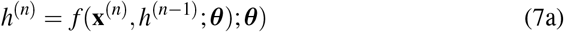

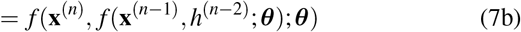

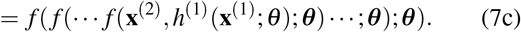

The idea behind sharing ***θ*** in RNN states is similar to CNNs: parameter sharing across different time points allows RNNs to generalize the model to sequences of variable lengths, and share statistical strengths at different positions in time^117,150^. Similar to FFNNs, RNNs learn by propagating the gradients of each hidden state’s inputs at discrete times. This process becomes more intuitive if we consider the outputs of hidden units at various time iterations as if they were the outputs of different neurons in a deep multi-layer network. However, due to the sequential nature of RNNs, the back-propagation of gradients shrinks or grows at each time step, causing the gradients to potentially vanish or blow up. This fact, and the inability to parallelize training at different hidden states (due to the sequential nature of RNNs) makes RNNs notoriously hard to train, specially for longer sequences^127,151^. However, when these issues are averted (via gradient clipping or other techniques), RNNs are powerful models and gain state-of-the-art capabilities in many domains, such natural language processing. The training challenges combined with the nature of scRNAseq data have resulted in fewer developments of RNNs for single-cell analysis. However, recently some studies have used RNNs and Long Short-Term Memory^152^ (a variant of RNNs) used for predicting cell types and cell motility (*e.g*. see Kimmel et al.^153^).

### D. Residual Neural Network (RestNet)

As mentioned above, deep RNNs may suffer from vanishing or exploding gradients. Such issues can also arise in other deep neural networks as well, where gradient information could diminish as the depth increases (though approaches such as Batch Normalization^154^ aim to help with gradient issues). One way to alleviate vanishing gradients in very deep networks is to allow gradient information from successive layers to pass through, helping with maintaining information propagation even as networks become deeper. ResNets^155^ achieve this by skip (or residual) connections that add the input to a block (a collection of sequential layers) to its output. For a FFNN, consider function *f* in Eq. (2). Using the same notation as in Eq. (2), ResNet’s inner layers take the form shown in Eq. (8):

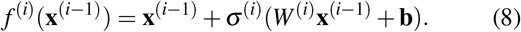

The addition of **x**^(*i*–1)^, the input of the current layer (or the output of (*i* – 1)-th layer), to the current i-th layer output is the *skip or residual* connection helps flow the information from the input deeper in the network, thus stabilizing training and avoiding vanishing gradient in many cases^155,156^. Indeed, this approach can be contextualized within the traditional time integration framework for dynamical system. For example, consider Eq. (9):

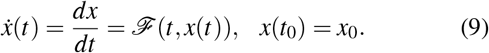

In the simplest case, this system can be discretized and advanced using *x*(*t_n_*) and some scaled value of 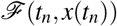, or a combination of scaled values of *ẋ*(*t_n_*). Forward Euler, perhaps the simplest time integrator, advances the solution as shown through the scheme in Eq. (10)

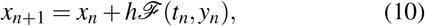

where *h* is a *sufficiently small* real positive value. ResNets use this idea to propose a different way of calculating the transformations in each layer, as shown in Eq. (8).

ResNets consist of residual blocks (also called modules), each of which containing a series of layers. For visual tasks, these blocks often consist of convolutional layers, followed by activation functions, with the skip connection adding the input information to the output of the residual blocks (as opposed to the individual layers inside). ResNets have different depths and architectures, with a number usually describing the depth of the model (*e.g*. ResNet50 means there are 50 layers [there are 48 convolution layers, one MaxPool and one AveragePool layers]).

ResNets have transformed DL by enabling the training of *very* deep neural networks, setting the state-of-the-art performance in many areas, particularly in computer vision^155^. The pre-trained ResNets on ImageNet dataset^157^ are widely used for transfer learning, where the network is either used as is or further fine-tuned on the specific dataset. Pre-trained ResNet models have also been used in spatial transcriptomics analysis, as we discuss later in this manuscript.

### E. Autoencoder (AE)

AEs^158,159^ are neural networks that aim to reconstruct (or copy) the original input via a *non-trivial mapping*. Conventional AEs have an “hour-glass” architecture (see Fig. 3(E)) consisting of two networks: (i) an encoder network, *Enc*(·), which maps an input 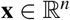 to a latent vector 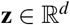 where, ideally, **z** contains the most important information from **x** in a reduced space (*i.e*. *d* ≪ *n*), (ii) the decoder network, *Dec*(·), which takes **z** as input and maps it back to 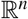, ideally, reconstructing **x** exactly; *i.e*. **x** = *AE*(**x**) = *Dec*(*Enc*(**x**)). AEs were traditionally used for dimensionality reduction and denoising, trained by minimizing a mean squared error (MSE) objective between the input data and the reconstructed samples (outputs of the decoder).

Over time, the AE framework has been generalized to stochastic mappings, *i.e*. probabilistic encoder-decoder mappings, *p_Enc_*(**z**|**x**) and *p_Dec_*(**x**|**z**). A well-known example of such generalization is Variational Autoencoders (VAEs)^160^, where by using the same hour-glass architecture, one can use probabilistic encoders and decoders to generate new samples drawn from an approximated posterior. Both traditional AEs and VAEs have practical applications in many biological fields, and have been used extensively in scRNAseq (see reference^10^ for an overview of these models), and are becoming more frequently employed in spatial transcriptomics analysis, which we overview later in this work.

### F. Variational Autoencoder (VAE)

One can describe VAEs^160^ as AEs that regularize the encoding distribution, enabling the model to generate new synthetic data. The general idea behind VAEs is to encode the inputs as a *distribution over the latent space*, as opposed to a single point (which is done by AEs). More specifically, VAEs draw samples **z** from an encoding distribution, *p_model_*(**z**), and subsequently feed the sample through a differentiable generator network, obtaining *Gen*(**z**). Then, **x** is sampled from a distribution *p_model_*(**x**;*Gen*(**z**)) = *p_model_*(**x**|**z**). Moreover, VAEs utilize an approximate inference network *q*(**z**|**x**) (*i.e*. the encoder) to obtain **z**. With this approach, *p_model_*(**x**|**z**) now is considered a decoder network, decoding **z** that comes from *q*(**z**|**x**). VAEs can take advantage of gradient-based optimization for training through *maximizing the variational lower bound*, 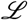, associated with **x**. Fig.3(F) depicts the architecture of traditional VAEs.

Mathematically, we can express the objective function as in Eq. (11):

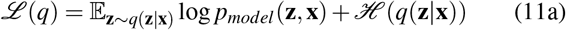

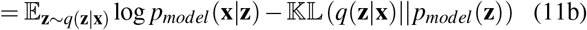

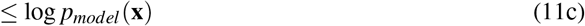

where 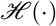 denotes entropy and 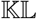 is the Kullback-Leibler divergence. The first term in Eq.(11c) is the joint loglikelihood of the hidden and visible variables under the approximate posteriors over the latent variables. The second term of Eq. (11c) is the entropy of the approximate posterior. This entropy term encorages the variational posterior to increase the probability mass on a range of **z** which could have produced **x**, as opposed to mapping to a one point estimate of the most likely value^117^.

Compared to other generative models (*e.g*. Generative Adversarial Networks (GANs)^161^), VAEs have desirable mathematical properties and training stability^117^. However, they suffer from two major weaknesses: (i) classic VAEs create “blurry” samples (those that adhere to an average of the data points), rather than the sharp samples that GANs generate due to GANs’ adversarial training. (ii) The other major issue with VAEs is posterior collapse: when the variational posterior and actual posterior are nearly identical to the prior (or collapse to the prior), which results in poor data generation quality^162^. To alleviate these issues, different algorithms have been developed, which have been shown to significantly improve the quality of data generation^163–168^. VAEs are used extensively for the analysis of single-cell RNA sequencing (see Erfanian et al.^10^), and we anticipate them to be applied to a wide range of spatial transcriptomics analysis as well.

## IV. DEEP LEARNING MODELS FOR SPATIALLY-RESOLVED TRANSCRIPTOMICS ANALYSIS

In the following sections, we describe the use of ML and DL to problems emerging from spatial transcriptomics.

### A. Spatial Reconstruction

Prior to the advancement of spatial transcriptomics, several studies aimed to reconstruct spatial information using gene expression data, with most of these works using a statistical framework. As perhaps one of the most influential models in this space, Satija *et al*.^59^ introduced Seurat: a tool which utilized spatial reference maps constructed from a few landmark *in situ* patterns to infer the spatial location of cells from corresponding gene expression profiles (*i.e*. scRNAseq data). This approach showed promising results: Satija *et al*. tested seurat’s capabilities and performance on developing zebrafish embryo dataset (containing 851 cells) and a reference atlas constructed from colorigenic *in situ* data for 47 genes^59^, confirming Seurat’s accuracy with several experimental assays. Additionally, they showed that Seurat can accurately identify and localize rare cell populations. Satija *et al*. also demonstrated that Seurat was a feasible computational solution for handling stochastic noise in omics data, and finding a correspondence between ST and scRNAseq data.

Although Seurat proved to be successful in some applications, it had the limitation of requiring spatial patterns of marker genes expression^60^. To alleviate Seurat’s limitations, newer methods that did not require spatial reference atlases were developed. A more recent and an influential model in this space is *novoSpaRc*^60^, with the ability to infer spatial mappings of single cells *de novo*. For novoSpaRc, Nitzan *et al*.^60^ assume that cells which are closer to one-another physically have similar gene expressions as well, therefore searching for spatial arrangement possibilities which place cells with similar expressions closer in space. Nitzan *et al*. formulate this search through a generalized optimal-transport problem for probabilistic embedding.

NovoSpaRc shows very promising results when it is applied to spatially reconstruct mammalian liver and intestinal epithelium, and embryos from fly and zebrafish from gene expression data^60^. However, novoSpaRc (and similar models) use a generic framework and can not be easily adapted to specific biological systems, which may be required given the vast diversity of biological processes and organisms. For this reason, many have utilized ML algorithms to specifically adapt to the biological system by learning from the data, as opposed to using pre-defined algorithms that remain unchanged. Indeed, we anticipate that DL models will soon play a salient role in spatial reconstruction of scRNAseq, given their ability to extract features from raw data while remaining flexible across different applications. In this section, we review DEEPsc^61^, a system-adaptive ML model which aims to impute spatial information onto non-spatial scRNAseq data.

**DEEPsc** is a spatial reconstruction method which requires a reference atlas (see Fig. 4). This reference map can be expressed as a matrix 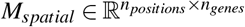 where 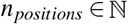 is number of spatial locations and 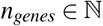 the number of genes. Maseda *et al*. start by selecting common genes between *M_spatial_* and the gene expression matrix, 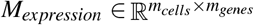 (where 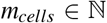 is the number of cells and 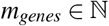 the number of genes), resulting in a spatial matrix 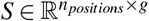 and an expression matrix 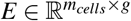, where 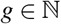 is the common genes between the two matrices. Next, *S* is projected into a lower dimension using principal component analysis (PCA), and the same PCA coefficients are used to project *E* into these principal components. In the last step of processing, both matrices are normalized by their largest elements, resulting all elements of the matrices *E* and *S* to be in [0, 1]. Let us denote the normalized and PCA-reduced spatial and gene expression matrices as 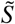 and *Ẽ*, respectively.

**FIG. 4.**
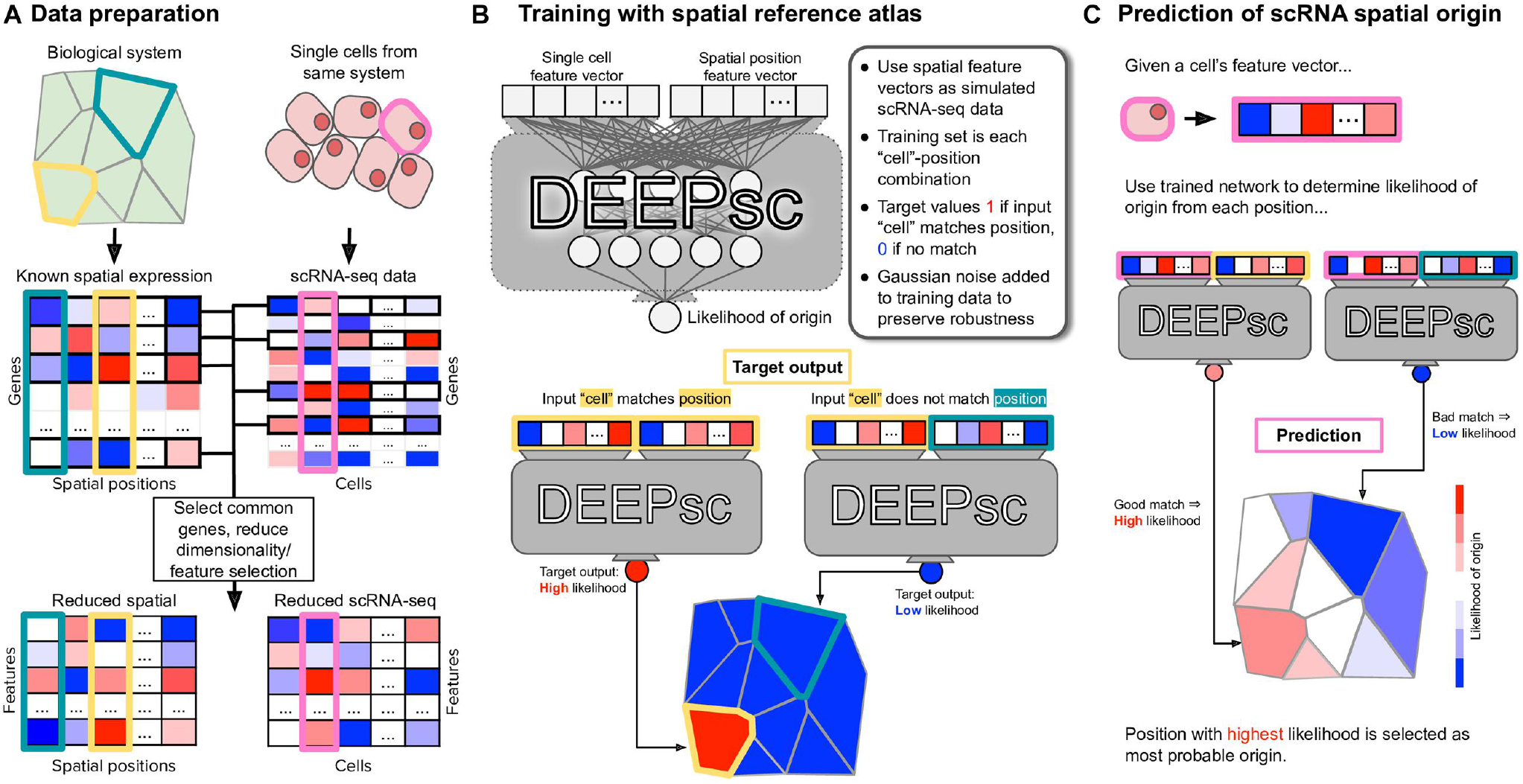
An overview of DEEPsc training and inference. **(A)** Maseda *et al*. find the common genes in both spatial and scRNAseq data, and perform dimensionality reduction on each data modality (with the final matrices having the same number of features). **(B)** During the training, DEEPsc uses spatial expression to “simulate” single-cell gene expression vectors. More specifically, every feature vector from the spatial expression is concatenated with all other vectors (labeled as “non-match”) and also itself (labeled as “match”) to form the input data to the neural network. **(C)** During inference, the scRNAseq feature vectors are concatenated with all spatial feature vectors, where the model should place a high probability for locations where the gene expression could have originated from. This figure was obtained from Maseda *et al*.^61^.

DEEPsc requires known spatial expression to learn the correct spatial positions, given the gene expression. More specifically, Maseda *et al*. construct training vectors of size 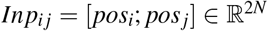 (with *N* being the number of features preserved in the reduced matrix 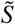). The first *N* elements of *Inp_ij_* correspond to the spatial expression at the *i*-th position, and the last *N* elements correspond to some position *j* in the reference atlas, including the position *j* = *i*. During training, DEEPsc’s goal is produce the highest likelihood when *j* = *i* (meaning that *Inp_ij_* = [*pos_i_*;*pos_i_*]), and assign low likelihood when *j* ≠ *i*. DEEPsc also adds Gaussian noise to *pos_i_* (the first *N* elements of *Inp_ij_*), which aims to preserve robustness and avoid overfitting. The addition of noise can lead to DEEPsc learning a complex nonlinear mapping between the spatial positions in the reference atlas rather than a simple step-like function which activates when an exact match is inputted. During inference stage (*i.e*. after DEEPsc is trained), *pos_i_* is replaces with the gene expression feature vector, which are the elements of *Ẽ*, and the goal is to predict the likelihood of the expression vector being originated from all possible positions *j*.

DEEPsc’s network is a FFNN with two hidden layers, with each 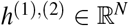, mapping to an *y* ∈ [0,1], where y can be viewed as a likelihood that the input cell originated from the input spatial position^169^. Given that for each training data *Inp_ij_* there will be *n_positions_* – 1 non-matches (labels of zero) and only one match for when *j* = *i*, the training labels will have many more zeros than ones. Therefore, Maseda *et al*. propose a non-traditional objective function which accounts for the imbalance between zeros and ones in the training data. This objective function is shown in Eq. (12)

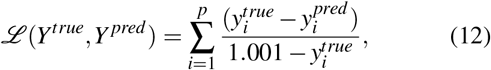

where 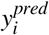 is the networks predicted outputs and 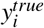 is the true target (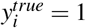 if it exactly matches, and 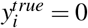 otherwise). This allows DEEPsc to avoid producing trivially zero outputs, which is important given the sparsity of the data. Maseda *et al*. also employ strategies in data splitting which helps to account for the inherent sparsity in the targets^61^. It is important to note that Maseda *et al*. also formulate a novel system-adaptive scoring scheme to evaluate the performance of DEEPsc using the spatial reference. However, the scoring scheme does not fall within the scope of this manuscript.

Maseda *et al*. apply DEEPsc for spatial imputation of four different biological systems (Zebrafish^59^, Drosophila^170^, Cortex^171^ and Follicle^172^), achieving accuracy comparable to several existing models while having higher precision and robustness^61^. DEEPsc also shows better consistency across the different biological systems tested, which can be attributed to its system-adaptive design. In addition, the authors attribute the performance and generalizibility of DEEPsc to the use of FFNN (which have been noted before in other biological applications as well^45,47^) and the various strategies for robustness used during the training of DEEPsc. On the other hand, a weakness of DEEPsc is its training time, which depends non-linearly on the number of locations available. However, this issue can be potentially mitigated by considering a small subset of possible locations, or a more optimized design when training the model.

### B. Alignment

*Alignment* in ST analysis refers to the process of mapping scRNAseq data to a physical domain while aiming to match the geometry with the available spatial data. As previously stated, NGS-based technologies suffer from limited capture rates and significant dropout^173^ (specially at higher resolutions). Before the use of DL in ST analysis, many computational approaches aimed to spatially reconstruct key marker genes scRNAseq data by assuming continuity in the gene space^60^, or by leveraging local alignment information^59^. Moreover, most techniques for alignment or deconvolution of spatial data either learned a program dictionary^32^ or estimated a probabilistic distribution of the data^64^ for the cell-types at each spot. However, such approaches are not generalizable to all experimental settings, since finding the mapping of sparse or sporadically distributed genes to the spots is difficult, and is error-prone due to dropouts^34^.

DL frameworks have the potential of providing robust models that can adapt to the specific species or technologies, while being generalizable to other datasets and platforms. The potential application of DL in alignment of spatially-resolved transcriptomics came to fruition recently through the work of Biancalani *et al*., called **Tangram**^34^. Tangram is a framework that, among many of its capabilities, can align scRNAseq or single-nucleus(sn) RNAseq profiles to spatial data; for the sake of simplicity, we refer to both data types as scRNAseq, although there are differences between the two methods (see reference^174^ for a systematic comparison of scRNAseq and snRNAseq approaches). Tangram aims to: (i) learn the transcriptome-wide spatial gene expression map at a single-resolution, and (ii) relate the spatial information back to histological and anatomical data obtained from the same samples. Tangram’s general workflow is to learn a mapping between the data modalities, and then to construct specific models for the downsteam tasks (such as deconvolution, correcting low-quality data, etcetera). We first summarize Tangram’s alignment algorithm, and then provide the applications in which DL models are utilized.

Tangram’s general objective is to learn a spatial matrix 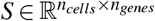 describing the spatial alignments for the cells, with *n_cells_*, *n_genes_* denoting the number of single-cells and number of genes, respectively. Let the expression of gene *k* in cell *i* be denoted by 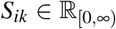, a non-negative value. Next, Tangram partitions (“voxelizes”) the spatial volume at the finest possible resolution (depending on the spatial technology) as a one-dimensional array. This allows Tangram to construct (1) a matrix 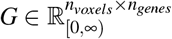 where *G_jk_* is a nonnegative value denoting the expression of gene *k* in voxel *j*, and (2) a cell-density vector **v** = {*v*_1_, *ν*_2_,…, *ν_n_voxels__*}, where 0 ≤ *ν_j_* ≤ 1 is the cell density in voxel *j* (with the total density for each voxel summing to 1).

The learning of transcriptome-wide spatial gene expression map at a single-resolution happens through learning a mapping operator 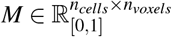 where *M_ij_* denotes the probability of cell *i* being in voxel *j*. Moreover, given any matrix 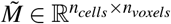, each element of the operator *M* is assigned according Eq. (13)

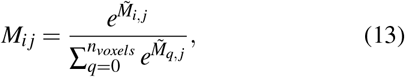

ensuring that 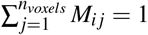, *i.e*. assigning a probability distribution along the voxels using the well-known *softmax*(·) function. Biancalani *et al*. define an additional quantity, *M_T_S*, which denotes the spatial gene expression as predicted by the operator *M*, and a vector **m** = {*m*_1_,…, *m_n_cells__*} where 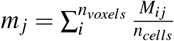 is the *predicted* cell density for each voxel *j*.

Given the preliminary quantities, we can now write Tangram’s generic objective function as shown in Eq. (14)

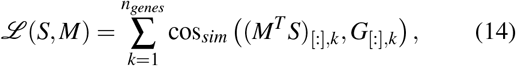

where “[:]” denotes the matrix slicing and cos_*sim*_ is the cosine similarity, defined as Eq. (15)

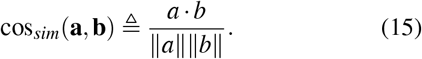

The objective function, Eq. (14), learns a proportional mapping of the genes to the voxels. Additionally, this loss function can be further modified to incorporate prior knowledge. Indeed, Biancalani *et al*. modify this to regularize for the learned density distributions and the cells contained within each voxel, as shown in Eq. (16)

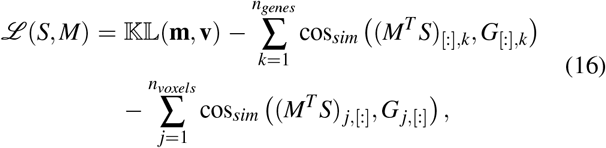

where minimizing the divergence (first term) enforces that the learned density distribution and the expected distribution are similar, and the additional loss over the voxels (third term) penalize the model if predicted gene expression is not proportional to the expected gene expression. Biancalani *et al*. minimize the objective shown in Eq. (16) through gradientbased optimizers implemented in PyTorch^175,176^. After optimizing Eq. (16), Tangram is able to map all scRNAseq profiles onto the physical space, thus performing alignment. It worthy to note that although Tangram learns a linear operator *M*, this mapping could be replaced with a deep neural network as well.

Tangram utilizes DL to integrate anatomical and molecular features, specifically for mouse brain images. To do so, the authors use an image segmentation network (U-Net^177^) in combination with a “Twin” network^178^ to produce segmentation masks of anatomical images, with both networks being CNNs. We present a general overview of these architectures in Fig. 5. The twin network uses DenseNet^179^: a deep NNs which concatenate the outputs at each layer to propagate salient information to deeper layers in the network (refer to section IIID for the motivation behind such approaches). More specifically, Tangram uses a pre-trained DenseNet encoder (trained on ImageNet) to encode images and remove technical noise and artifacts. Biancalani *et al*. also add two additional layers to the pre-trained encoder, which map the outputs to a smaller latent space. The encoder of the twin network is fine-tuned on learning the prediction of spatial depth difference between two images: Two random images are inputted to the twin network with their spatial difference depth being the desired target output, *d^true^*. The network then tries to predict the depth, *d^pred^*, for all *N* inputs, ultimately comparing them against the corresponding true depth differences, *d^true^*(as shown in Eq. (17))

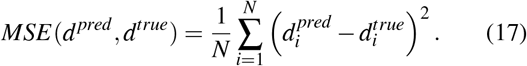

**FIG. 5.**
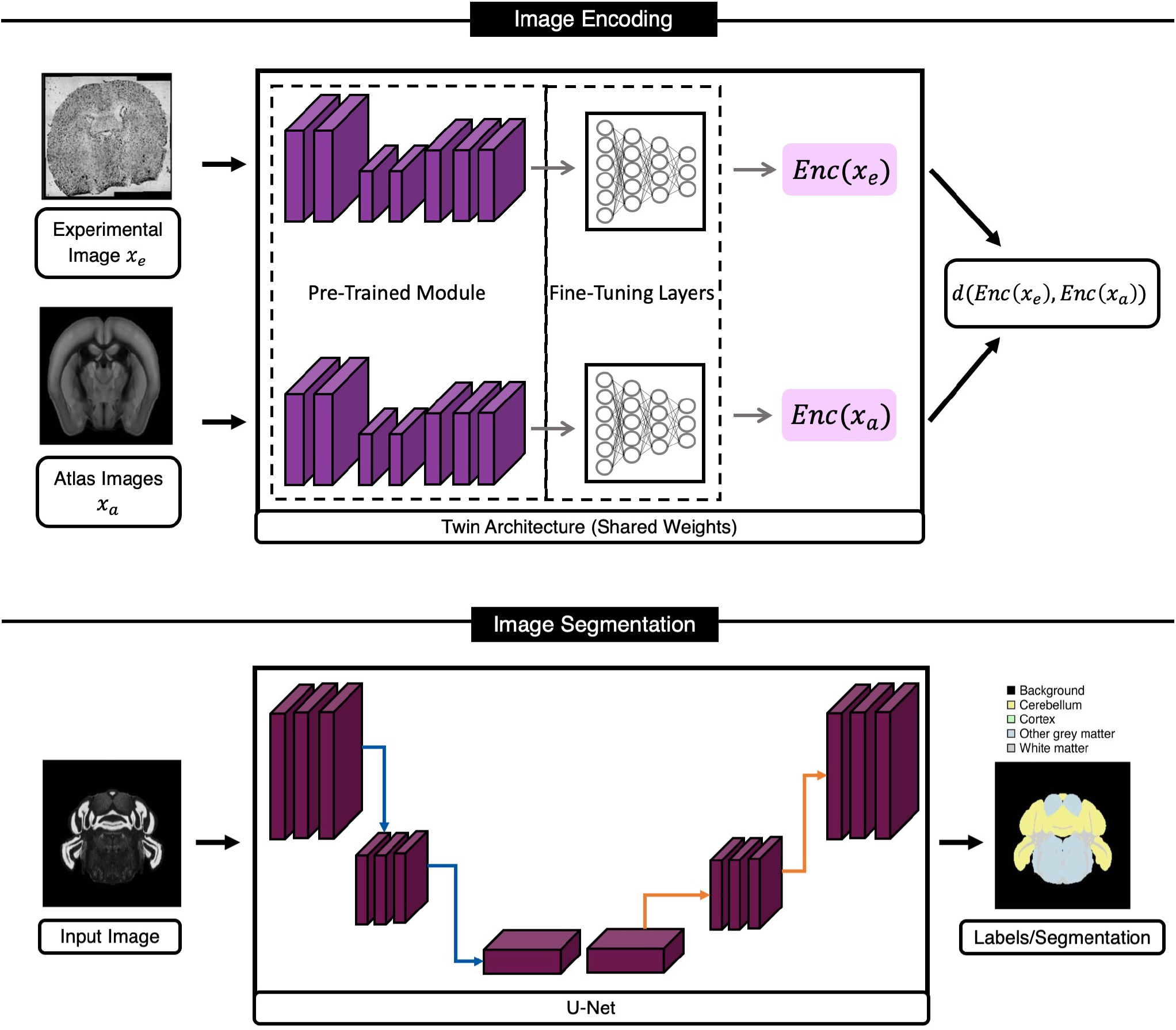
Tangram’s DL Framework for Aligning/Integrating Histology and Anatomical Data with Molecular Data. Tangram’s model for this task is a combination of encoding (using a twin network) and segmentation modules. The twin network learns a similarity metric for brain sections based on anatomical features in images, while the U-Net model is trained to segment five different classes on mouse brain images. This figure was recreated for this manuscript using images from Biancalani *et al*.^34^.

The segmentation model of Tangram generates five custom segmentation masks (background, cortex, cerebellum, white matter, and other gray matter) which are compatible with existing Allen ontology atlas. The segmentation model is a U-Net, which uses a pre-trained ResNet50^155^ as its core. Finally for each pixel in input images, the model’s last layer (a softmax function) assign a probability of belonging to one of the five segmentation classes. Tangram’s segmentation model aims to optimize the superposition of the cross entropy and Jaccard index, as presented in Eq. (18)

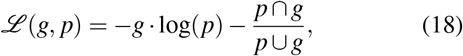

with *p* denoting the model prediction and *g* referring to the ground truth image.

Biancalani *et al*. demonstrate that Tangram learns an accurate mapping between the spatial data and scRNAseq gene expression when applied to fine or coarse grained spatial atlases. The authors show that their approach works well across different technologies (namely ISH, smFISH, Visium, STARmap and MERFISH) at different resolutions and gene coverage, and is able to learn a robust and accurate alignment mapping for the isocortex of the adult healthy mouse brain^34^. While Tangram can offer different advantages based on the spatial technology, it can produce consistent spatial mappings and overcoming limitations in resolution or throughput, which is beneficial in many ST experiments and studies.

### C. Spot Deconvolution

One downside of using NGS-based technologies remains to be their resolution: Despite the recent technological advancements, most ST platforms (*e.g. Spatial Transcriptomics*, Visium, DBiT-seq^180^, Nanostring GeoMx^181^ and SlideSeq) do not have a single-cell resolution. The number of cells captured in each spot still varies based on the tissues (about 1-10^182^) and the technology used. On the other hand, we can not assume that all cells within a spot are the same, due to the heterogeneity of the cells. Therefore, it is necessary to use computational approaches for inferring the cell types in each spot or voxel. Such estimations would be possible if there were a complementary scRNAseq dataset. The process of inferring the cellular composition of each spot is known as *cell-type deconvolution*. Deconvolution has been at the forefront of computational efforts and it is important in building oragan atlases^20,183,184^. In fact, cell-type deconvolution is an existing procedure for inferring cell-type composition in RNAseq data using scRNAseq. However, methods developed for bulk RNAseq do not account for the spatial components of ST datasets, and are therefore generally inadequate. Given that deconvolution is an existing practice in RNAseq studies, we will refer to spatial deconvolution problem as *spot deconvolution* to distinguish between the traditional methods and the ones developed for ST analysis.

We divide spot deconvolution methods into three categories: (i) Statistical methods, (ii) Machine Learning and (iii) Deep Learning, with many of the current models falling into the first two categories. We now dive deeper into the two existing models which use DL for performing spot deconvolution.

### D. DestVI

DestVI (DEconvolution of Spatial Transcriptomics profiles using Variation Inference) is a Baysian model for spot deconvolution. DestVI employs a conditional deep generative model (similar to scVI^185^, a popular model for scRNAseq analysis) to learn cell-type profiles and continuous sub-celltype variations, aiming to recover the cell-type frequency and the average transcriptions state of cell-types at each spot. To do so, DestVI takes a pair of transcriptomics datasets as inputs: (i) a reference scRNA-seq data and (ii) a query spatial transcriptomics data (from the same samples). DestVI then outputs the expected proportion of cell types for every spot and a continuous estimation of cell-state for the cell types present in each spots, which can be viewed as the average state the cell-types in each spot, which Lopez *et al*. suggest as useful for downstream analysis and formulation of biological hypotheses^69^.

DestVI uses two different latent variable models (LVMs) for distinguishing cell-type proportions and delineating celltype-specific sub-states (shown in Fig. 6). The first LVM is for single-cell data (therefore named *scLVM*) which assumes the counts follow a negative binomial (NB) distribution, which has shown to model RNAseq count data well^185–187^. Specifically, Lopez *et al*. assume that for each gene *g* and cell *n*, the count of observed transcripts, *x_ng_*, follows a NB distribution paramaterized with (*r_ng_, p_g_*): *r_ng_* = *l_n_* · *f*(*γ_n_, c_n_*; ***θ***) is a parameter which depends on the type assigned to the cell *c_n_*, the total number of detected molecules *l_n_*, and a low-dimensional latent vector *γ_n_* (which lopez *et al*. set *γ_n_* = 5) that describes the variability of cell-type assignment to cell *c_n_*, and a neural network *f* parameters ***θ*** (in this case, a two layer NN). The second parameter of the NB, *p_g_*, is optimized using variational Bayesian inference. We can summarize the assumptions for scLVM as shown in Eq. (19):

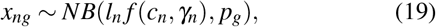

with the latent variable 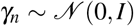. Each *c_n_* (the annotations) are represented by a one-hot encoded vector, which is concatenated with *γ* to serve as the input of the NN *f*. Lopez *et al*. use a VAE to optimize for the marginal conditional likelihood log *p_θ_* (*x_n_*|*l_n_, c_n_*).

**FIG. 6.**
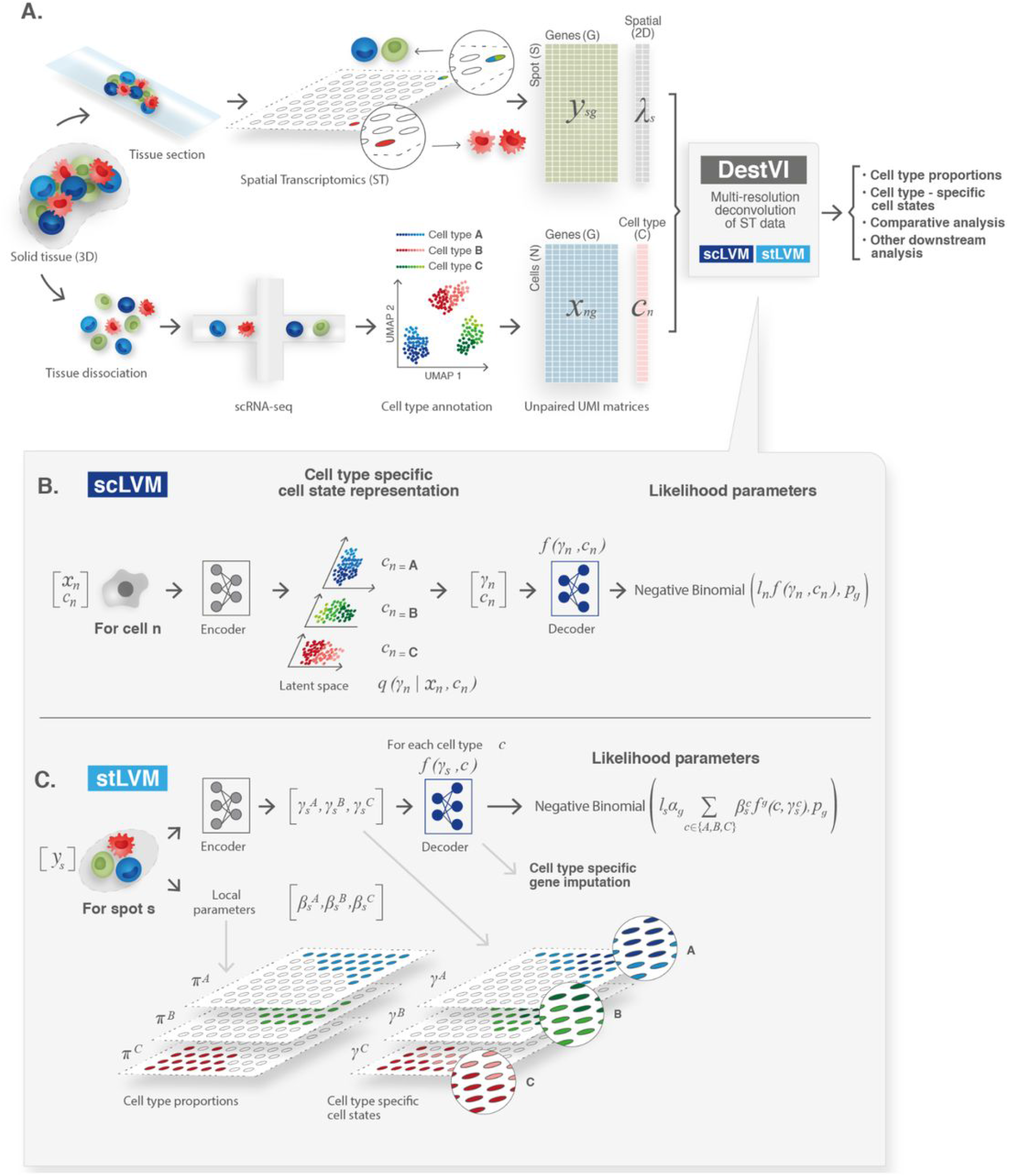
Visualization of DestVI’s Computation Workflow for Spot Deconvolution. DestVI uses information from both data modalities of ST data (shown in **A**). DestVI defines two latent variable models (LVMs) for each data modality: an LVM for modeling scRNAseq data (**B**) and one that aims to model the ST data (**C**). We describe each one in section IV C. This image is re-used from Lopez *et al*.^69^

Finally for scLVM, a mean-field Gaussian distribution *q_ϕ_*(*γ*|*c_n_, x_n_*), parametrized by another two-layer NN *g*, is inferred for each cell which quantifies the cell state and the associated uncertainty. The NN *g* takes a concatenation of (i) the gene expression vector *x_n_* and (ii) the one-hot encoded cell annotations as its inputs. The network *g* outputs the mean and variance of the variational distribution for *γ_n_*, obtained through optimizing Eq. (20)

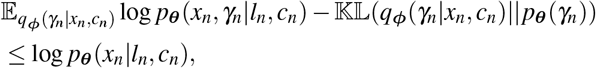

where *p*_θ_(*γ_n_*) is the prior likelihood for *γ_n_*. Similar to training any other VAE, the observations are split in mini-batches and sampling from the variational distribution is done using the reparameterization trick described in Kingma *et al*.^160^. The computational workflow for scLVM is visualized in Fig. 6(B).

The second LVM aims to model the spatial transcriptomics data (hence called *stLVM*) with the assumption that the number of observed transcripts *x_sg_* at each spot *s* for each gene *g* follow a NB distribution. Additionally, Lopez *et al*. also assume that each spot has *C*(*s*) cells, with each cell *n* in spot *s* being generated from the latent variables (*c_ns_, γ_ns_*). For stLVM’s NB distribution, the rate parameter *r_sg_* = *α_g_l_s_f_g_*(*c_ns_, γ_ns_*;***θ***_*g*_), where *α_g_* is a correction factor for the gene-specific bias between spatial and scRNAseq data, *l_g_* is the overall number of molecules observed in each spot, and *f_g_* is a NN network with parameters ***θ***_*g*_. These assumptions and quantities allow Lopez *et al*. formulate the total gene expression *x_sg_* as shown in Eq. (21)

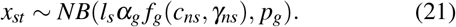

Moreover, using a parameter to designate the abundance of every cell type in every spot, *β_sc_*, and NB’s rate-shape parameterization property (see Aragón et al.^188^), Eq. (21) can be rewritten as in Eq. (22)

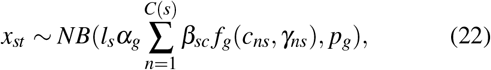

supposing that cells from a given cell type *c* in a spot *s* must come from the same covariate 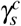.

The covariate 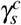 in DestVI allows for the model to account for ST technology discrepancies by assuming various empirical priors (refer to Fig. 6). Lopez *et al*. simplify the problem of identifying every cell type in each spot to determining the density cell-types, *i.e*. assuming that there cannot be significantly different cell states of the same cell types within a spot. Lopez *et al*. use a penalized likelihood method to infer point estimates for *γ^c^, α*, and *β*. With the addition two strategies to stablize the training of DestVI, the final objective function for stLVM consists of (i) the negative binomial likelihood (ii) the likelihood of the empirical prior and (iii) the variance penalization for *α*.

Lopez *et al*. use simulations to present DestVI’s ability to provide higher resolution compared to the existing methods and estimate gene expression by every cell type in all spots. Furthermore, they show that DestVI is able to accurately deconvolute spatial organization when applied mouse tumor model. In the cases tested, Lopez *et al*. demonstrate that DestVI is capable of identifying important cell-type-specific changes in gene expression between different tissue regions or between conditions, and that it can provide a high resolution and accurate spatial characterization of the cellular organization of tissues.

### E. DSTG

Deconvoluting spatial transcriptomics data through graphbased convolutional network (DSTG)^65^ is a recent semisupervised model which employs graph convolutional networks (GCN)^134^ for spot deconvolution. DSTG utilizes scRNAseq to construct a pseudo-ST data, and then building a link graph which represents the similarity between all spots in both real and pseudo ST data. The pseudo-ST is generated by combining scRNAseq transcriptomics of multiple cells to mimic the expression profiles at each spot; while the real ST data is unlabeled, the pseudo-ST has labels. To construct the link graph, DSTG first reduces the dimensionality of both real and pseudo data using canonical correlation analysis^189^, and then identifies mutual nearest neighbors^190^. Next, a GCN is used on the link graph to propagate the real and pseudo ST data into a latent space that is turned into a probability distribution of the cell compositions for each spot.

Song *et al*. form the link graph by taking the number of spots as the number of vertices, resulting in a graph *G* = (*V, E*) where |*V*| denotes the number of spots and *E* represents the edges between them. DSTG takes two inputs: (i) the adjacency matrix of graph *G*, represented by *A*, and (ii) a combination of both real and pseudo datasets *X* = [*x_pseudo_*; *x_real_*] ∈ ℝ^*m*×*N*^ where *m* is the number of variable genes, and *N* = *S_pseudo_* + *S_real_* (the total number of spots in both datasets) with *S_pseudo_* and *S_real_* indicating the number of spots in the pseudo and real ST datasets, respectively. Next, Song *et al*. normalize the adjacency matrix (for efficient training of DSTG) using the diagonal degree of *A*, denoted by *D*, as shown in Eq. (23)

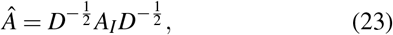

where *A_I_* = *A* + *1* (with *I* denoting the identity matrix). Given the two inputs, DSTG’s graph convolution layers take the following form:

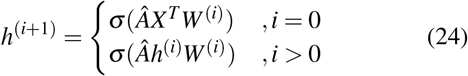

with *h*^(*i*)^ denoting the *i*-th hidden layer of DSTG, and *σ*(·) being a non-linear activation function (in this case *σ*(·) = *ReLU*(·)). The output of DSTG is denoted by *y_s,t_*, the proportion of cell type *t* = {0,···, *T*} at each each spot s = {0, ···, *N*}. Song *et al*. design DSTG’s architecture as shown in (25):

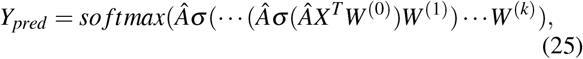

where *k* is the last layer, and 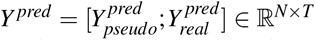 is the predicted proportions at each spot in the pseudo and real data, denoted by *Y_pseudo_* and *Y_real_*, respectively. It is important to note that Song *et al*. chose a GCN with three layers after performing an ablation study on the number of layers. Finally, DSTG is trained by optimizing the cross entropy loss:

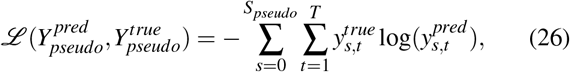

with 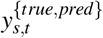 denoting the label for true/predicted cell type t at spot *s*. Note that this constitutes a semi-supervised training for DSTG, since only labels for the pseudo ST are used in training, but the model will also learn to predict labels for the real dataset as well (refer to Eq. (25)).

Song *et al*. note that, compared to traditional approaches, DSTG provides three key advantages: (i) Given that DSTG uses variable genes and a non-linear GCN, it allows for learning complex deconvolution mappings from ST data. (ii) The weights assigned to the different cell types in the pseudo-ST and the semi-supervised scheme allow DSTG identify key features which allow the model to learn the cellular composition in real data. (iii) DSTG’s scalability and adaptability will be beneficial in ST analysis, given that the sequence depth of ST data is expected to increase. Song *et al*. show that DSTG consistently outperforms the benchmarked state-of-the-art model (SPOTlight) on both synthetic data and real data. More specifically, DSTG is evaluated on simulated data generated from PBMC where it shows high accuracy between the predicted cell compositions and the true proportions. Song *et al*. also find that DSTG’s deconvolution of ST data from complex tissues including mouse cortex, hippocampus, and human pancreatic tumor slices is consistent with the underlying cellular mixtures^65^.

### F. Spatial Clustering

Clustering allows the aggregation of data into subpopulations based on some shared metric of distance or “closeness”. In RNA sequencing studies, clustering is the first step of identifying cell clusters, often followed by laborious manual annotation (*e.g*. through identifying differentially expressed genes) or some automated workflows^191^. Clustering has been a crucial step in many scRNAseq studies, often performed using graph-based community detection algorithms (such as Louvain^192^ or Leiden^193^) or more traditional methods (such as K-Means^118^). Although the scRNAseq techniques can be used in some ST studies (*e.g*. for multiplexed FISH data where single-cell resolution is available), the result may be discontiguous or erroneous since the spatial coordinates have not been taken into account^74^. Therefore, there is a need for ST-specific methods that can utilize both gene expression and histology data to produce clusters that are coherent, both in gene expression and physical space.

Recently, new frameworks for spatial clustering of ST data have emerged which utilize both spatial and expression information available. Zhu *et al*.^71^ introduced a Hidden-Markov Random Field (HMRF)-based method to model the spatial dependency of gene expression using both the sequencing and imaging-based single-cell transcriptomic profiling technologies. HMRF is a graph-based model used to model the spatial distribution of signals. Using the ST data, Zhu *et al*. create a grid where neighboring nodes are connected to each other. However, the spatial pattern can not be observed directly (since it is “hidden”), and it must be inferred through observations that depend on the hidden states probabilisticly. Similar to Zhu *et al*., BayesSpace^73^ employs a Bayesian formulation of HMRF, and uses the Markov chain Monte Carlo (MCMC) algorithm to estimate the model parameters. Despite the ability of these methods to cluster voxels (or cells) into distinct subpopulations, these approaches suffer from the lack of versatility required to handle different modalities present in ST data^74^.

With the emergence of newer technologies, the scale and variability within datasets are increasing, requiring more general and flexible models for accurate and robust analysis of these studies. A few ML-based approaches have been proposed to combat some of these mentioned challenges. Below, we review the ML-based methods which offer scalability and are generally more applicable to various experimental settings.

### G. SpaCell

SpaCell^72^ is a double-stream DL framework which utilizes both histology images and the associated spot gene counts. For the histology data, Tan *et al*. first preprocess the images (removing low-quality images, stain normalization, normalizing the pixels using a z-transform and removing background noise). Next, they split each histology image into tiles that contain one spot each (sub-images of 299 × 299 pixels). For the preprocessing of the count matrix (which contains the reads at each spot), Tan *et al*. follow traditional scRNAseq preprocessing workflows, including count normalization, removal of outlier genes and cells with too few genes. After the preprocessing stage, each tile (containing the image of a spot) corresponds to a column in the count matrix (reads from the same spot). At this point, each image 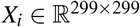 and the count matrix *M* will be in a 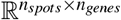 space. However, Tan *et al*. reduce the count matrix to only contain 2048 most variable genes at each spot, therefore resulting in a new count matrix 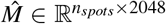. Let us denote the *i*^th^ spot of 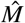 as 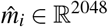, which has a corresponding image *x_i_*.

In order to spatially cluster cells of the same type, both image and count data must be used. The first step in SpaCell is to pass on the spot images, *x_i_*, to a pre-trained ResNet50 (trained on ImageNet data) in order to output feature vectors, 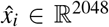 (each having the same dimensionality as columns of M). Next, to extract features from both modalities, SpaCell uses two separate AEs for the image feature vectors, 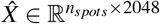, and the most-variable-genes counts, M, with both AEs having the same latent dimension (we discuss the reason behind this later). Let us denote the AE for images as *AE_I_*(·) = *Dec_I_*(*Enc_I_*(·)), and the gene counts AE as *AE_G_*(·) = *Dec_G_*(*Enc_G_*(·)), with *Enc*_{*I,G*}_(·) and *Dec*_{*I,G*}_(·) indicating the encoder and decoders, respectively.

Given *N* spots, each AE in spaCell aims to minimize three objective functions for their respective inputs [i.e. 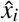 for *AE_I_*(·) and 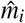 for *AE_G_*(·)]: (i) the mean squared error (MSE) between the input and output (shown in Eq. (27)), (ii) the KL divergence between the probability distributions for input and constructed output of all N spots (denoted by *p* and *q* in Eq. (28) respectively) and (iii) Binary Cross Entropy (BCE), shown in Eq. (29):

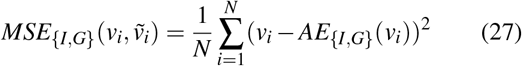

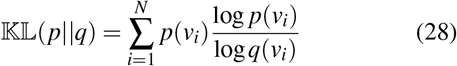

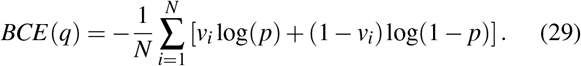

Once training has concluded, spaCell encodes both images and gene counts, *i.e*. 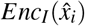 and 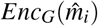, to be used for clustering. More specifically, clustering is performed on a matrix that is the concatenation of the latent vectors produced by each AE, 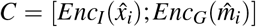. This is why the latent spaces of *AE_I,G_*(·) have the same dimension. After obtaining the concatenated matrix, the downstream clustering is performed using K-Means (which can be substituted for other algorithms as well). Through this procedure, spaCell uses both data modalities and can produce clusters that are highly accurate when compared to the true clusters (annotated by pathologists).

### H. SpaGCN

SpaGCN^74^ is a graph convolution network (GCN) that integrates both spatial information and histology images to perform spatial clustering. Using each spot as vertices, Hu *et al*. create a weighted undirected graph, *G* = (*V, E*), where |*V*| is the total number of spots and *E* is the set of edges with prescribed weights representing the similarity between the nodes. The weight of each of these edges is determined by (i) the distance between the two spots (nodes) that the edge connects, and (ii) the associated histology information (in this case, pixel intensity). This means that two spots are deemed similar if they are physically close to one another *and* they seem similar in the histology image.

In order to attribute the pixel information to each spot, Hu *et al*. use mean RGB pixel intensity of each spot within a window of size 50 × 50 pixels. That is, given a spot *s* with physical coordinates (*x_s_, y_s_*) and pixel coordinates (*x_ps_,y_ps_*), SpaGCN calculates the mean and variance of all the pixels present within a 50 × 50 pixels centered at (*x_ps_,y_ps_*). Let *ps_r_, ps_g_, ps_b_* denote the means, and *var_r_*(*ps*), *var_g_*(*ps*), *var_b_*(*ps*) refer to the variance for the red, green and blue channels, respectively. SpaGAN then summarizes the pixel mean and variance information as a unified value, as shown in Eq. (30)

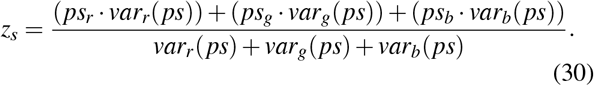

Furthermore, *z_s_* is rescaled using the mean and standard deviation of each coordinate (including the newly-created *z* axis), with an additional scaling factor which can put more emphasis on histology data when needed. Let *μ_z_* denote the mean of *z_s_*, and let *σ_x,y,z_* be the standard deviation of *x_s_, y_s_, z_s_* with *s* ∈ *V*, then we can formulate the rescaling as the following:

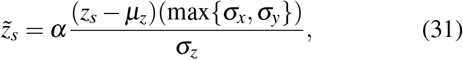

where *α* denotes the scaling factor described previously (*α* = 1 by default).

Using the rescaled value in Eq. (31), the weight of each edge between two vertices *s* and *k* is calculated as shown in Eq. (32)

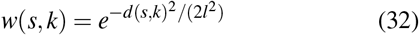

where *l* denotes the characteristic length scale and *d*(*s, k*) is the traditional Euclidean distance, as shown in (33)

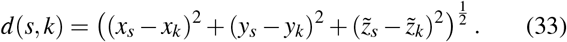

SpaGCN’s network construction (and backpropogation) is similar to other GCNs, inspired by Kipf *et al*.^134^ (for an overview of GCNs, refer to section IV E). The network intakes the adjacency matrix *A* to represent the graph *G*, and a reduced-dimension representation of the gene expression matrix, which Hu *et al*. achieve using PCA with 50 principal components. The outputs of the GCN network is matrix which includes combined information on histology, gene expression, spatial position. SpaGCN then uses the output of the GCN to perform unsupervised clustering of the spatial data.

SpaGCN uses the Louvain algorithm (an iterative unsupervised clustering algorith) on the output of GCN to initialize cluster centroids, with the number of clusters (controlled by Louvain’s *resolution* parameter) being optimized on maximizing the Silhouette score^194^). The iterative updates are based on optimizing a metric that defines the distance between each spot and all cluster centroids using the t-distribution as a kernel. For a centroid *c_j_*, a total of *N* clusters, and the embedded point *h_i_* for spot *i*, this metric can be defined as the probability of assigning cell *i* to cluster *j*, as shown in Eq. (34)

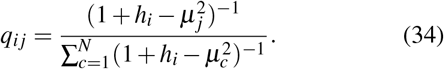

Hu *et al*. further refine the clusters using an auxiliary target distribution (shown in Eq. (35)) which prefers spots assignments with the highest confidence, and normalizes the centroid contribution to the overall loss function as the following:

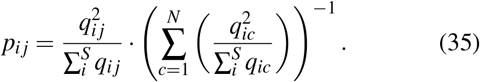

Lastly, spaGCN is trained by optimizing the 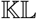 divergence between the *p* and *q* distributions, as shown in Eq. (36)

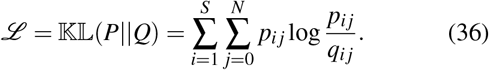

Hu *et al*. demonstrate that SpaGCN can accurately identify spatial clusters that are consistent with manual annotations, since SpaGCN utilizes information from both gene expression and histology. The authors perform spatial clustering with SpaGCN on human dorsolateral prefrontal cortex, and human primary pancreatic cancer and multiple mouse tissue data, showing that SpaGCN performs consistently well, outperforming other state-of-the-art models (stLearn, BayesSpace, and Louvain). These results show the feasibility and potential of SpaGCN for clustering spatial-resolved transcriptomics.

### I. Cell-Cell Interactions

Multicellular organisms depend on intricate cell–cell interactions (CCIs) which dictate cellular development, homeostasis, and single-cell functions^195^. Unravelling such interaction within tissues can present unique insights on complex biological processes and disease pathogenesis^195–198^. CCI has been investigated using both scRNAseq and RNAseq, wherein most approaches test for enrichment in ligand-receptor profiles in the expression data^199–201^. However, ST data can offer a more comprehensive view of CCI, since the distance traveled by ligand signal is crucial in determining the type of cell–cell signaling^182^. Given the importance of CCI and the advantages that ST data provides, several computational approaches for inferring cellular interactions using ST data have been developed, such as SpaOTsc^78^, Giotto^81^, MISTy^80^.

SpaOTsc is a model that can be used in integrating scRNAseq data with spatial measurements, and in inferring cellular interactions in spatial-resolved transcriptomics data. SpaOTsc aims to estimate cellular interactions by analyzing the relationships between ligand-receptor pairs and their downstream genes. SpaOTsc formulates a spatial metric using the optimal transport algorithm, returning a mapping that contains the probability distribution of each scRNA-seq cell over a spatial region. SpaOTsc also utilizes a random forest in order to infer the spatial range of ligand-receptor signaling and subsequently removing the long-distance connections. Another approach is Giotto^81^: Giotto is an extended and comprehensive toolbox designed for ST analysis and visualization, which includes a CCI model which calculates an enrichment score (the weighted mean expression of a ligand and the corresponding receptor in the two neighboring cells). Giotto then constructs an empirical null distribution by moving the locations for each cell-type, subsequently calculating corresponding statistical significance (P-value), and ordering the ligand-receptors pair-wise for all neighboring cells.

Although the mentioned models have shown to discover simple cellular interactions, such approaches often fail to detect complex gene-gene interactions, which is essential in understanding many diseases. DL models learn such complicated interactions from raw data, further utilizing ST data in studying CCI. For this purpose, **StLearn**^79^ is a recent DL model that, among many of its capabilities, can learn CCI from spatially-resolved transcriptomics. StLearn’s DL components lies within its Spatial Morphological gene Expression (SME) normalization. The SME normalization aims to combine critical information from Hematoxylin and Eosin stained (H&E) tissue images and transcriptome-wide gene expression profile to then take advantage of in downstream analysis, such as clustering, spatial trajectory inference, and CCI.

The SME normalization procedure includes *(i) spatial location:* In order to use the spatial positions for selecting neighboring spot pairs, Pham *et al*. consider two spots *s_i_* and *s_j_* as neighbors if the center-to-center euclidean distance between two spots,|*C*(*s_i_*) – *C*(*s_j_*)|, is less than a specified distance *r, i.e*. |*C*(*s_i_*) – *C*(*s_j_*)|< *r*. Pham *et al*. include all paired spots *s_i_* and *s_j_* as input to adjust for the gene expression of the center spot *s_i_*. The next step in SME normalization is *(ii) Morphological similarity:* stLearn calculates the morphological similarity between spots using feature vectors produces by an ImageNet-pre-trained ResNet50. More specifically, all H&E images corresponding to each spot *s_i_* is inputted to the ResNet50 model, which then produces a feature vector 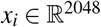. Subsequently, stLearn performs PCA on each feature vector *x_i_*, resulting in reduced-dimension feature vectors 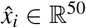. To calculate the morphological distance (MD) between two neighboring spots *s_i_* and *s_j_* (according to criterion defined in *(i)*), Pham *et al*. measure the cosine similarity between two reduced feature vectors (refer to Eq. (15) for definition of Cosine Similarity); this MD is shown in Eq. (37),

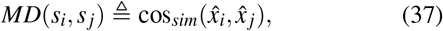

As a last step in SME normalization, the gene expression at each spot *s_i_* is normalized using the MD distance, as shown in Eq.(38)

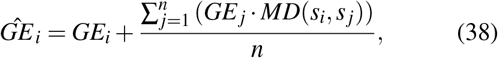

where *GE_i_* denotes raw gene expression counts at spot *s_i_*, and *n* is the total number of neighbors identified for spot *s_i_*.

After SME normalization, stLearn can perform multiple downstream tasks, including the identification of tissue regions with high CCI activities^79^. StLearn’s CCI algorithm finds ligand–receptor (L-R) co-expression between neighboring spots, and tests for the enrichment of L-R pairs between two cell types, which are compared to a random null distribution using CellPhoneDB^200^. After this initial testing, significant L-R pairs are selected to calculate cell-cell interactions. These interactions are measured using the nearest neighbors for a spot, and are queried to the cKDTree algorithm^202^ to validate that the neighboring cells express ligand or receptor genes that are above a pre-defined threshold. Next, Pham *et al*. form a matrix where significant L-R pairs represent the features (columns) for each spot coordinates (the rows). Using this matrix, stLearn can cluster the spatial regions with the most similar L-R co-expression values, which combined with the CCI measures, can identify tissue regions that have high L-R co-expression, indicating areas that have a high likelihood of active CCI. This approach consitutes stLearn as one of the the first methods which combines both spatial cell populations (identified through clustering) and L-R interactions to detect tissue region with a high likelihood of CCI. Pham *et al*. apply stLearn’s CCI method to breast cancer tissue and identify spatial regions and L-R pairs in cancer-immune cell interactions, indicating a great potentials for shedding light on CCI using ST data.

## V. CONCLUSIONS AND OUTLOOK

The ST field is rapidly growing, with new datasets and analysis pipelines released weekly. The innovations in biological methods will continue to spur the creativity in algorithm development, with an emphasis on ML-based frameworks. Although the space of DL models for ST analysis is currently small, we anticipate the field to experience a paradigm shift towards deep-learned models. In this review, our goal was to provide readers with the necessary biological, mathematical, and computational background for understanding the existing approaches, and expanding upon the current models to address the challenges posed by the ST domain.

In this manuscript, we provided an overview of current DL-based techniques for alignment and integration of ST data, spatial clustering, spot deconvolution, inferring cell-cell communication, and approaches for reconstructing spatial coordinates using scRNAseq data (with limited or no spatial reference atlas). The DL methods we presented, in comparison to their conventional counterparts, offer accuracy and scalability advantages. However, DL methods are not always the preferred choice as they are computationally expensive and may lack biological interpretability. As more methods for ST analysis are developed, we believe that standard datasets for benchmarking new models as well as comprehensive accuracy and efficiency analysis of existing techniques will be of significant value to the field. Though the existing methods set the new state-of-the-art in their respective categories, room for improvements remains large. Among the ST downstream analyses, applications of DL algorithms for studying cell-cell communication and identification of spatially-variable genes remain mostly underexplored. Given DL models’ ability to extract sophisticated patterns from raw data, we anticipate that DL approaches will prove useful in unraveling complex biological processes, aiding the efforts in identifying cellular interactions and highly variable genes in a spatial context.

Recent technological advancements have enabled researchers to utilize various single-cell omics sources to construct multi-omics datasets, providing comprehensive view of many diseases (*e.g*. COVID19^203,204^ and cancer^205^), and developmental processes^206,207^. As the single-cell analysis enters the multi-omics age, the need for integrating ST data with other single-cell sources will increase. Therefore, we expect an increase in ML-based frameworks for data integration and alignment, spearheaded by DL-based approaches. Additionally, due to the noise and multi-modality of ST data, there exists an unmet need for methods that account for batch effects in spatial and gene expression data. Given the success of DL techniques for batch effect removal in scRNAseq, we foresee DL models being widely used for batch effect correction of spatially-resolved transcriptomics data.

Despite the recentness of ST technologies, researchers have successfully used these technologies to generate spatially-resolved cell atlases, providing new insights on a wide range of biological processes and organs^208–212^. Such studies show the tremendous potential that ST technologies hold, but also highlight the need for scalable and efficient analyses tools. The application of DL to ST analysis remains a rapidly evolving nascent domain, demonstrating promising great prospects in advancing the field of ST, and the integration of ST datasets with other omics data.

## ACKNOWLEDGMENTS

We wish to acknowledge Oscar Davalos for offering fruitful discussions and providing useful insights on earlier versions of this work. We also would like to thank Tommaso Buvoli, Maia Powell, and the anonymous reviewers for providing valuable feedback on earlier drafts of this manuscript. The authors received support from the National Institutes of Health (R15-HL146779 and R01-GM126548) and the National Science Foundation (DMS-1840265).

## GRAPHICS ACKNOWLEDGMENTS

The Visium slide and its visualizations in Fig. 1 and 3 were accessed from 10x Genomics^213^. The mouse brain histology image in Fig. 1 was taken from reference^214^. The illustration of mouse brain in Fig. 2 was obtained from BioRender^215^.

## AUTHOR CONTRIBUTIONS

A.A.H. and S.S.S. wrote the manuscript, edited the drafts and conceptualized the figures, which A.A.H then created.

